# Incentive Salience, not Psychomotor Sensitization or Tolerance, Drives Escalation of Cocaine Self-Administration in Heterogeneous Stock Rats

**DOI:** 10.1101/2025.10.04.680105

**Authors:** Jarryd Ramborger, Joseph Mosquera, Molly Brennan, Benjamin Sichel, Dyar Othman, Sonja Plasil, Elizabeth Sneddon, Selene Zahedi, Alex Morgan, Supakorn Chonwattanagul, Kathleen Bai, Lindsey China, Tu La, Lisa Maturin, Lieselot L. G. Carrette, Olivier George

## Abstract

Sensitization and tolerance are two phenomena often studied independently despite overlapping neurobiological substrates. Each has extensive research showing their influence on the development and maintenance of addiction, but the degree to which they drive escalation in cocaine self-administration is poorly understood. Using self-administration, intravenous noncontingent infusions, and pose-estimation machine vision, we find that incentive salience, not psychomotor sensitization or tolerance, drives the escalation of cocaine self-administration in heterogenous stock rats. Individual differences in psychomotor sensitization or tolerance were found to have no effect on cocaine intake. Incentive salience as measured by locomotion and active lever entrances per meter traveled occurring before the self-administration session began (pre-lever activity measures) during Short Access (2-hours) was found to predict intake during Long Access (6-hours). Both pre-lever locomotion and active lever entrances per meter were found to increase during Long Access and after two-to-three days of abstinence. Critically, rats with low pre-lever activity during Short Access escalated both their intake and pre-lever measures by the end of Long Access to levels comparable with high pre-lever activity rats who maintained their elevated responding. These findings support the notion that incentive salience during Short Access is a catalyst to escalated use and an early marker of addiction vulnerability. Moreover, they suggest that individuals initially resistant to incentive salience can, with sufficient exposure, become sensitized and escalate cocaine use to the same level as more susceptible individuals. Analysis of pre-lever activity offers a novel longitudinal behavioral marker to predict vulnerability and provides a framework for understanding individual trajectories of addiction.

## 1. Introduction

Addiction is a chronic, relapsing disorder characterized by cycles of binge/intoxication, withdrawal/negative affect, and preoccupation/anticipation, leading to long-lasting neuroadaptations in stress, executive function, and reward processing [1,2]. Within this framework, two of the most studied adaptations to repeated drug exposure are tolerance, a reduced response to the drug, and sensitization, an enhanced response. Both processes have been demonstrated in preclinical models of addiction, but the expression of either phenomenon varies widely depending on experimental parameters such as drug of choice, method of exposure (e.g., noncontingent injections versus self-administration), schedule of access (e.g. Short Access, Long Access, Intermittent, etc.), and outcome variables (e.g. locomotion, physiological measures, and operant behavior) [3–13]. Despite evidence that tolerance and sensitization share overlapping neurobiological substrates, they are often investigated in isolation [14–17], leading to two competing models in the transition to addiction.

According to the hedonic homeostatic dysregulation theory, 1) repeated drug use leads to tolerance to the positive reinforcing effects of drugs, and 2) the allostatic recruitment of stress circuitry, thereby shifting the hedonic set point and motivating continued use to restore homeostasis [1,18]. In contrast, the incentive-sensitization theory proposes that repeated use hypersensitizes neural circuits leading to 1) *incentive salience*—drug-related cues become increasingly "wanting" even as “liking" diminishes—and 2) *behavioral sensitization*—a heightened psychomotor response observed primarily in psychostimulants [19]. Both phenomena can contribute to the addiction cycle; however, the degree to which sensitization and tolerance drive escalation of cocaine self-administration is poorly understood. While several studies have demonstrated the expression of tolerance or sensitization to cocaine with noncontingent methods of continuous, repeated, or single infusions [7,9,20–23], or the use of different lengths of self-administration access [3–6,24–26], none have in a longitudinal paradigm employing noncontingent challenges, Short Access, and Long Access.

To address this gap, using an operant cocaine self-administration paradigm in conjunction with machine vision pose-estimation, we assessed multiple behavioral indices to differentiate individual development of tolerance or sensitization and the subsequent influence on cocaine self-administration. These included: 1) locomotion following a single noncontingent intravenous infusion of cocaine at three time points (see **Fig. 1A** for timeline); 2) locomotion for 15 minutes before the self-administration session begins (pre-lever); 3) drug-seeking during pre-lever and Noncontingent sessions, measured as nose entrances into the active lever area per meter traveled; 4) cocaine intake within the first 15 minutes and 5) full session; and 6) breakpoint on a progressive ratio schedule. Psychomotor tolerance and sensitization will be assessed by locomotion during Noncontingent sessions. Incentive salience will be assessed by both locomotion and nose entrances into the active lever area per meter during the pre-lever phase of self-administration.

**Figure 1.**
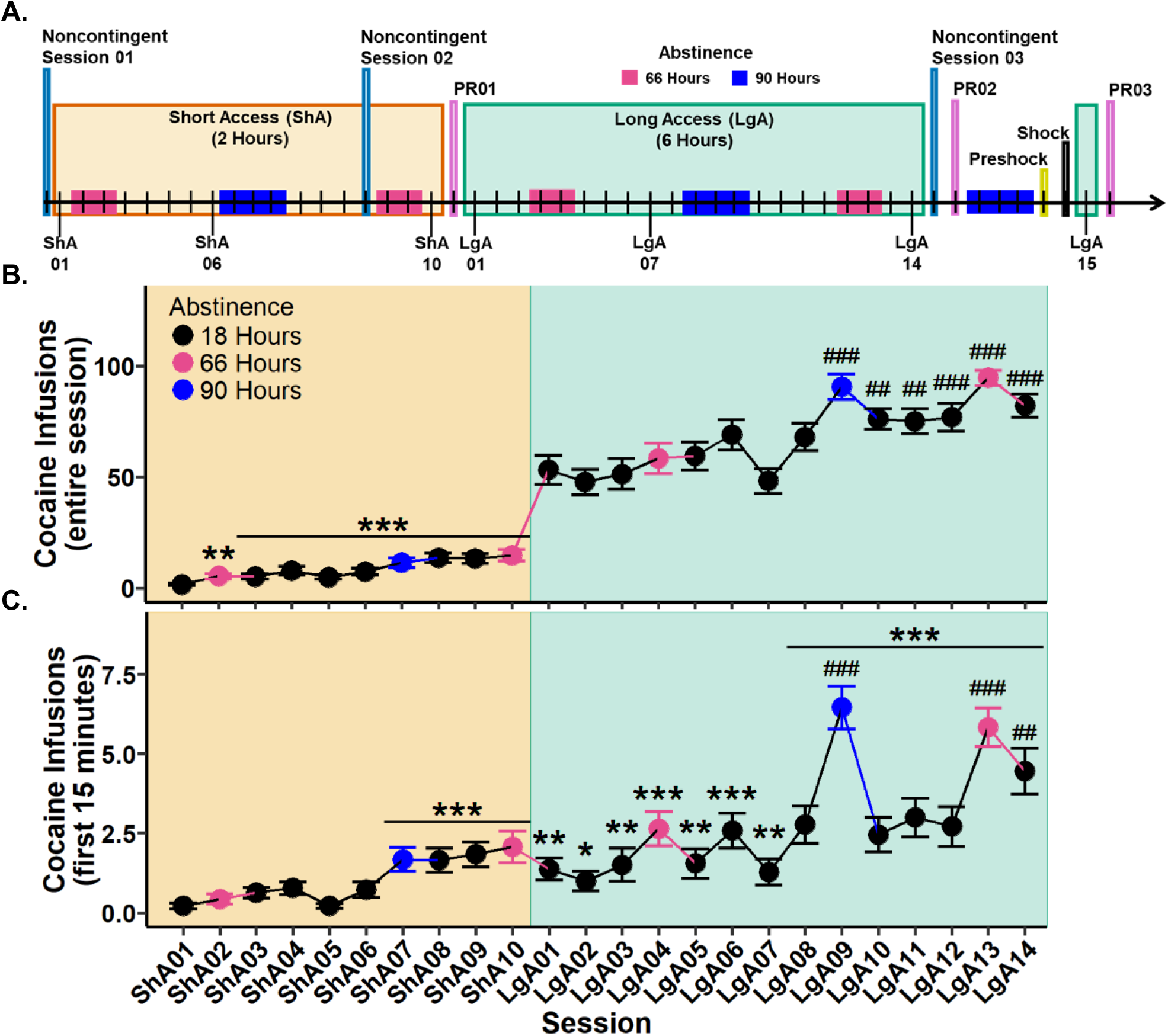
Experimental timeline and cocaine infusions during Short (ShA) and Long Access (LgA). **(A.)** Experimental timeline with 66 hours (pink) and 90 hours (blue) indicating instances of abstinence. **(B.)** Cocaine infusions during the entire session. **(C.)** Cocaine infusions during the first 15 minutes of the session. Plots represent means and SEM. \**p*<.05, \*\**p*<.01, \*\*\**p*<.001 vs. Short Access 01; ##*p*<.01, ##*p*<.001 vs. Long Access 01.

## 2. Materials and Methods

Procedures in sections 2.1-2.3 below have been published elsewhere in more detail [27–28]. ChatGPT4 was utilized for coding assistance and authors take full responsibility for output.

### 2.1. Subjects

Heterogeneous stock rats (N=37; n=21 males; n=16 females) obtained from the UC San Diego colony [29] were used to mimic human genetic diversity and housed in pairs on a 12-hour light/dark cycle. Rats were anesthetized with 1–5% isoflurane for jugular vein catheterization using Micro-Renathane tubing (Braintree Scientific) connected to a guide cannula that exits through the back and secured with mesh and dental acrylic. Post-surgery, rats received flunixin subcutaneously (2.5 mg/kg) and cefazolin intramuscularly (330 mg/kg). Catheters were flushed daily with heparinized saline (10 U/ml, American Pharmaceutical Partners) mixed with 0.9% bacteriostatic sodium chloride (Hospira) and cefazolin (52.4 mg/0.2 ml). Catheter patency was tested intravenously with brevital sodium (1.5 mg, American Pharmaceutical Partners) at experiment start and end, with failing rats removed from the experiment. All procedures were conducted in strict adherence to the National Institutes of Health Guide for the Care and Use of Laboratory Animals and were approved by the Institutional Animal Care and Use Committees of UC San Diego.

### 2.2. Drugs

Cocaine HCl (National Institute on Drug Abuse, Bethesda, MD) was dissolved in 0.9% sterile saline. During self-administration sessions, a press of the active lever resulted in an intravenous infusion of 0.1ml at 0.5 mg/kg over six seconds by pump, followed by a 20-second timeout where the active lever had no programmed effect. During Noncontingent sessions, a single intravenous infusion of 0.3ml at 1.5 mg/kg (i.e. three times the dose received in self-administration) was delivered in under five seconds in the same operant chamber as self-administration to address influences of environment and intravenous infusion rate on the development of sensitization [30–33].

### 2.3. Self-Administration

Over 8 weeks (Monday-Friday), rats participated in 30 self-administration sessions of various lengths and ratios. Under fixed ratio one, rats participated in 10 two-hour Short Access sessions,15 six-hour Long Access sessions, one one-hour Preshock, and one one-hour Shock session. During Shock, each infusion had a 30% chance to be paired with a footshock (0.3 mA for 0.5 sec) through the grid floor. Under progressive ratio, rats participated in three six-hour sessions in which the response requirements for an infusion followed the equation [5e(injection number × 0.2)] – 5. The breakpoint was defined as the last ratio before a 60-minute period in which a ratio was not completed, in which case the session ended. A complete experimental timeline is shown in **Fig. 1A**.

### 2.4. Noncontingent Sessions

Seven days after surgeries, rats underwent the first noncontingent session (Noncontingent 01) which occurred before Short Access 01. Rats were first video recorded for 30 minutes to establish baseline measurements (Baseline 01), then given an intravenous saline injection and recorded for 30 minutes (Saline 01), followed by a noncontingent intravenous injection of cocaine (1.5 mg/kg) and recorded for 30 minutes (Drug 01). The same procedures were replicated for the subsequent noncontingent sessions.

### 2.5. Video Recording and Processing

#### 2.5.1. Video Recording

Video recordings were conducted as described previously [34] with a few alterations. A static FFmpeg build (ffmpeg-6.0-armhf-static; https://johnvansickle.com/ffmpeg/) was installed onto each Raspberry Pi for video processing to reduce video size threefold without sacrifice to video quality. All sessions were recorded for 30 minutes. During self-administration sessions, levers were extended into the chamber after 15 minutes of recording. Each comma-separated value (csv) file was processed in R version 4.3.1 using RStudio [35–36] to ensure consistent length and alignment to the lever extension, resulting in ∼25 minutes per csv, or ∼12.5 minutes for both pre- and post-lever data. Out of 1,110 csv files (37 rats over 30 sessions), 26 were excluded for various issues such as catheter or infrared light disconnections. For eight rats during Long Access 04 the Med Associates program was initiated too early, resulting in no pre-lever data. For Noncontingent sessions, post-processing resulted in ∼25 minutes per csv. Out of 333 csv files, six files failed to track during Social LEAP Estimates Animal Poses (SLEAP) processing.

#### 2.5.2. SLEAP Modeling

A pose-estimation model was trained using SLEAP on ∼5200 labeled frames over 89 videos utilizing the top-down configuration and a unet backbone [37]. Thirteen nodes were tracked: nose, ears, neck, four paws, base (where tail met the body), active and inactive levers, active lever cue light, and the catheter node, which was chosen as the model anchor point. A second model was trained to track Noncontingent 01 as rats were mistakenly not plugged into their catheter lines. This model focused on six nodes to accelerate training: nose, ears, neck, base, and catheter as the anchor node, utilizing ∼2200 labeled frames from 40 videos.

#### 2.5.3. High Throughput Computing on Open Science Grid

Video recordings were processed on the OSG Consortium supercomputer system [38–41]. A python script with SLEAP command line functions processed each video to identify one instance (track) using an “instance” identification method and Hungarian approach between frames. Results of each node for each frame inference registered as (*x*, *y*) coordinates were interpolated to fill missing data points (https://sleap.ai/notebooks/Analysis_examples.html).

### 2.6. Dependent and Independent Variables

Statistical calculations were conducted using RStudio. Euclidian distances calculated movement of the catheter (locomotion) and the nose (nose motion) nodes frame-by-frame and converted from pixels to centimeters using Image2 [42] and the known measurements of the ENV-005 grid. Given the barrel distortion of a fisheye lens, the conversion factor was computed by averaging two length measurements (left and right) and three width measurements (top, middle, and bottom), resulting in ∼0.111cm/px. Locomotion variables are the total distances traveled smoothed by a window of 10 (10 FPS, i.e., 0.33 seconds) and converted to meters. Locomotion (m) pre-lever is the distance traveled during the 15 minutes before the levers are extended into the chamber. Active and inactive lever entrances were calculated by producing a bounding box around the lever nodes (*x*,*y*) at 30 +/- *x* and 18 +/- *y*, and counting each entrance of the nose node into the lever areas. Active and inactive lever entrances per meter are the total entrances normalized by locomotion. Cocaine infusions are the total infusions delivered during the session, and cocaine infusions first 15 minutes are those that occurred during the video recordings. Session corresponds to an individual session (e.g. Short Access 01). Session type refers to experimental phase (e.g. Long Access). Abstinence refers to the time between the end of one session and the beginning of the next, where 18 Hours translates to consecutive days, 66 Hours to a two-day weekend, and 90 Hours to a three-day weekend. Pre-lever variables were examined by abstinence to test for effects that could facilitate sensitization [6,14–17,20,30–34,43–44]. Analyses with sex as a factor were performed and reported in the Supplementary Material as most effects were not sex dependent.

### 2.7. Statistical Analyses

Statistical comparisons were made by fitting a linear mixed-effects model utilizing the lmerTest package [45–46] or a generalized linear mixed-effects model utilizing the glmmTMB package [47] in R. Given that linear and generalized linear models are inherently robust to moderate deviations from normality and extreme values, outliers for all dependent variables were winsorized at the 95th percentile to minimize their influence while retaining potentially meaningful behavioral variability. Inferential tests of fixed effects for linear models were conducted using Type III ANOVA with Satterthwaite degrees of freedom approximations (Type III ANOVA test) which have been shown to control for Type I errors in hierarchical models [48], while generalized linear models a Type III Wald chi-square analysis of deviance was conducted (Type III Wald test). Model assumption tests and fit were analyzed with the simulateResdiduals() function from the DHARMa package [49]. The model fitting process and chosen models of each analysis can be found in the Supplementary Material with best adherence to suggestions previously published [50] but prioritizing the results of DHARMa tests. Transformations of response variables where necessary (see Supplemental Material) were achieved using Box-Cox or Yeo-Johnson after identifying the most effective lambda using powerTransform() from the car package [51]. Post hoc analyses were conducted using contrasts(), while direct baseline comparisons used test(), both within the emmeans package and calculate the pairwise comparisons of the estimated marginal means and are therefore model-derived ratios or mean differences [52]. All *p*-values were adjusted by the Benjamini-Yekutieli (BY) method due to its consideration of dependence and correlations in nested data such as mixed models [53]. From the rstatix package, Student’s t-tests were conducted with t_test() with Bonferroni adjustments, and correlations with cor_test() by either Pearson or Spearman Rank as determined by a normality test with shapiro_test() [54]. Plots represent the raw average values or percent difference from baseline.

## 3. Results

### 3.1. Cocaine Infusions

#### 3.1.1. Cocaine infusions escalate during both Short and Long Access

To determine whether cocaine intake changed during Short Access, a Type III Wald test was performed. A significant main effect of session was observed (*χ*²(9)=100.94, *p*<.001). Post hoc analyses revealed all sessions to have significantly more cocaine infusions in comparison to Short Access 01. These results confirm that cocaine intake increased as early as Short Access 02 (**Fig. 1B**).

To determine whether cocaine intake changed during Long Access, a Type III Wald test was performed. A significant main effect of session was identified (*χ*²(13)=173.01, *p*<.001). Post hoc comparisons indicated that Long Access 09 through Long Access 14 had significantly more infusions relative to Long Access 01. These results show that cocaine intake continued to escalate into the third week of Long Access (**Fig. 1B**).

#### 3.1.2. Cocaine infusions during the first 15 minutes escalate across sessions

To evaluate whether cocaine infusions during the initial 15 minutes of each session escalated, a Type III Wald test was conducted. A significant main effect of session was identified (*χ*²(23)=304.87, *p*<.001). Post hoc analyses revealed that cocaine infusions were significantly greater from Short Access 07 through Long Access 14 compared to Short Access 01. Additionally, when compared to Long Access 01, only sessions Long Access 09, 13, and 14 showed significantly elevated infusions (**Fig. 1C**). These results indicate that escalation of cocaine intake during the first 15 minutes emerged at the end of Short Access and continued throughout Long Access.

### 3.2. Locomotion During Noncontingent Sessions

#### 3.2.1. Noncontingent cocaine administration increases locomotion with no evidence of psychomotor sensitization or tolerance following Short or Long Access cocaine self-administration

To determine if raw locomotion differs between Noncontingent sessions following noncontingent administration of cocaine, a Type III ANOVA test was performed. A significant main effect of session was found (*F*(8,290.88)=17.38, *p*<.001). Post hoc analyses only found significant differences within each respective Noncontingent session. Specifically, Drug sessions had significantly greater locomotion than their respective Baseline and Saline sessions (**Fig. 2A**).

**Figure 2.**
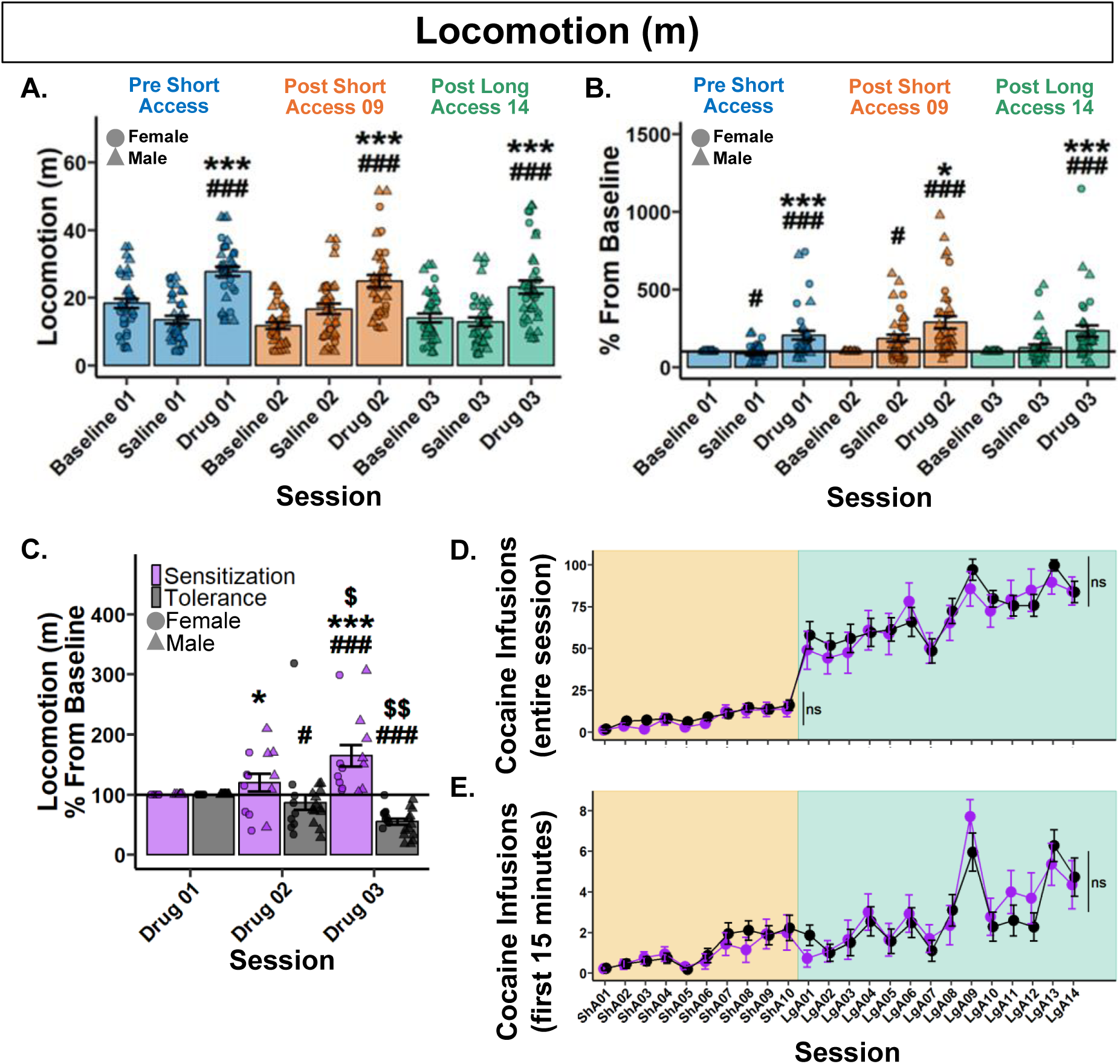
Psychomotor sensitization and tolerance show no effect on cocaine self-administration. **(A.)** Raw locomotion (m) across Noncontingent sessions. *###p*<.001 vs Baseline; \*\*\**p*<.001 vs Saline. **(B.)** Locomotion (m) percent difference from Baseline. #*p<*.05, *###p*<.001 vs Baseline; \**p*<.05, \*\*\**p*<.001 vs Saline. **(C.)** Locomotion (m) percentage difference from Baseline (Drug 01) grouped by behavioral expression. Drug 01 < Drug 03, Sensitization (n = 14); Drug 01 > Drug 03, Tolerance (n = 22). *#p*<.05, *###p*<.001 vs Baseline; Within behavioral expression groups: $*p*<.05, $$*p*<.01 vs Drug 02; Between behavioral expression groups: \**p*<.05 vs Tolerance Drug 02, \*\*\**p*<.001 vs Tolerance Drug 03. **(D-E.)** Cocaine infusions over the entire session (D.) and within the first 15 minutes (E.) grouped by behavioral expression. Plots represent means and SEM.

To assess the change in the locomotor response to noncontingent cocaine administration, data was analyzed as percent difference from each rat’s respective Baseline session. A Type III ANOVA revealed a significant main effect of session (*F*(5,176.97)=16.36, *p*<.001). Post hoc analyses revealed that Drug sessions were significantly greater than their respective Saline and Baseline sessions, but no significant differences between Drug sessions (**Fig. 2B**). These results show that locomotion did not differ between Noncontingent sessions following noncontingent administration of cocaine at the population level.

#### 3.2.2. Rats differentially develop psychomotor sensitization or tolerance after a history of cocaine self-administration

To assess each individual rat’s response to noncontingent infusions, rats were grouped by their locomotion difference from Drug 01 to Drug 03. Rats who *increased* were labeled “Sensitization” and rats who *decreased* were labeled as “Tolerance”.

Locomotion between Drug sessions was then analyzed by a percentage change from baseline (Drug 01). Following a 0.13 lambda Box-Cox power transformation, a Type III Wald test found no significant main effect of behavioral expression (*χ*²(1)=4.8, *p*=.059), but a significant effect of session (*χ*²(1)=6.47, *p*=.031) and their interaction (*χ*²(1)=15.2, *p*<.001). Post hoc analyses revealed that Tolerant rats significantly decreased in locomotion in Drug 02, with a further decrease in Drug 03. Conversely, Sensitized rats increased in Drug 03 and were greater than Tolerant rats in both Drug 02 and 03. These results demonstrate divergent profiles in the psychomotor response to cocaine that begins after just nine Short Access sessions and continues through Long Access (**Fig. 2C**). The effect observed was not due to stereotypies in Tolerant rats (**Fig. S1**).

#### 3.2.3. Psychomotor sensitization and tolerance show no effect on cocaine self-administration

To test whether the development of psychomotor sensitization or tolerance influenced cocaine self-administration, a Type III Wald test was performed for both Short and Long Access. During Short Access, a main effect of session was found (*χ*²(9)=64.36, *p*<.001), but not behavioral expression (*χ*²(1)=0.98, *p*=.967), or their interaction (*χ*²(9)=8.72, *p*=.967. Similarly, in Long Access, a main effect of session was found (*χ*²(13)=107.44, *p*<.001), but not behavioral expression (*χ*²(1)=0.47, *p*=1), or their interaction (*χ*²(13)=11.54, *p*=1). These results suggest that the development of psychomotor sensitization or tolerance to the effect of cocaine has no effect on cocaine intake during Short or Long Access (**Fig. 2D**).

To determine if the development of psychomotor sensitization or tolerance influences cocaine infusions in the first 15 minutes, a Type III Wald test was performed. A main effect of session was found (*χ*²(23)=154.13, *p*<.001), but not behavioral expression (*χ*²(1)=0.003, *p*=1), or their interaction (*χ*²(23)=30.78, *p*=.356). These results confirm that the development of psychomotor sensitization or tolerance to the effect of cocaine has no effect on loading behavior during cocaine self-administration (**Fig. 2E**).

### 3.3. Pre-Lever Activity

#### 3.3.1. Locomotion pre-lever increases from Short to Long Access and predicts cocaine intake during the first 15 minutes of self-administration

To determine if locomotion pre-lever increased over the experiment, a Type III Wald test was performed. A significant main effect of session was found following a 0.29 lambda Box-Cox power transformation (*χ*²(23)=204.7, *p*<.001). Post hoc analyses revealed significant increases at Short Access 07, Short Access 08, Long Access 09, and Long Access 13 in comparison with Short Access 01 (**Fig. 3A**). These results indicate that locomotion pre-lever increased from Short to Long Access.

**Figure 3.**
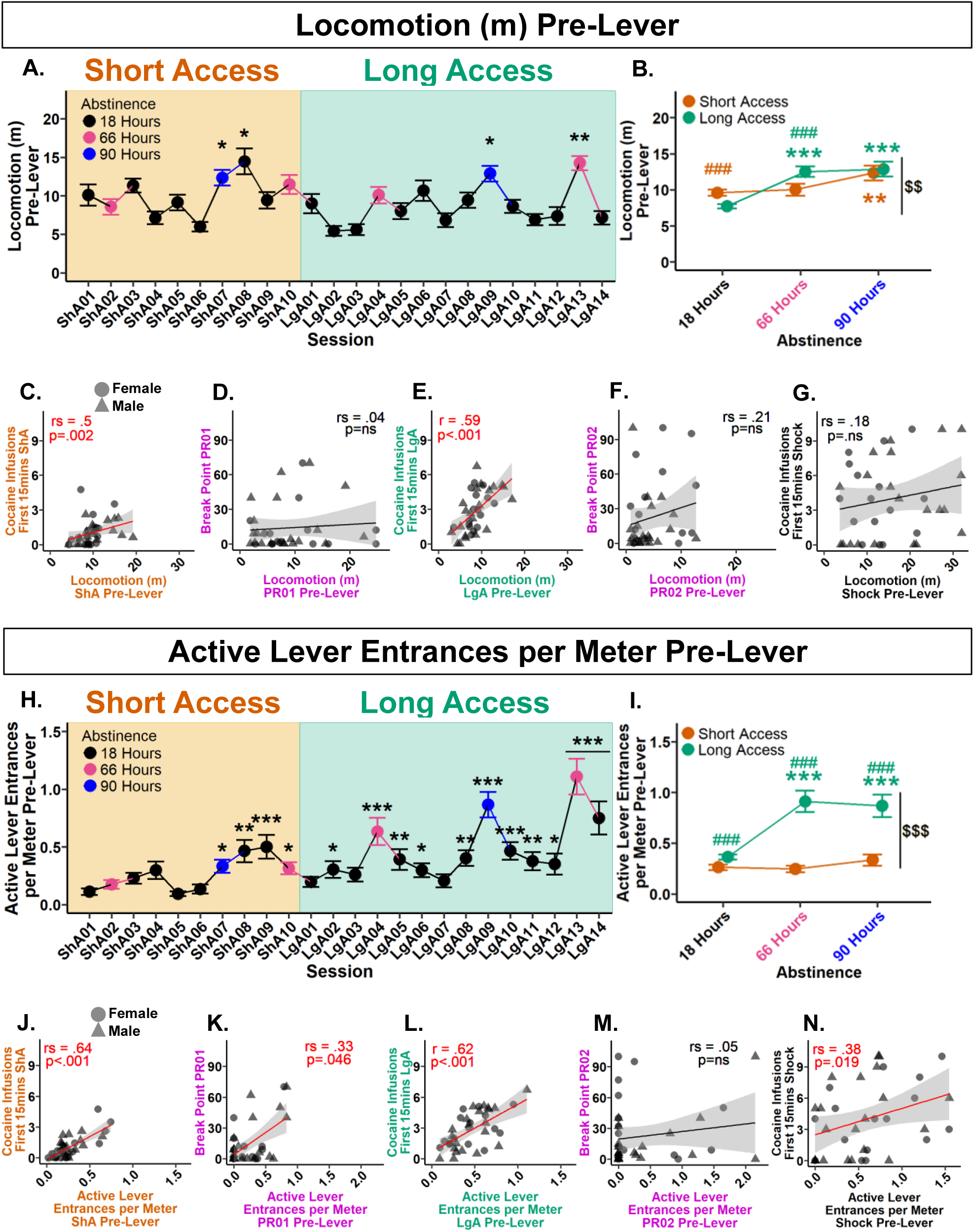
Pre-lever activity increases with Long Access, hours of abstinence, and is predictive of cocaine infusions within the first 15 minutes of self-administration. **(A.)** Locomotion pre-lever across experiment. \**p*<.05, \*\**p*<.01 vs Short Access 01. **(B.)** Comparison of abstinence levels within and between session types. Within session type: \*\**p*<.01, \*\*\**p*<.001 vs 18 hours; between session type and abstinence: ###*p*<.001 vs 18 hours Long Access, ###*p*<.001 vs 66 hours Short Access; between session type: $$*p*<.01 vs Short Access. (C.) Mean locomotion pre-lever during Short Access correlates with mean cocaine infusions during first 15 minutes of Short Access. **(D.)** Locomotion pre-lever during Progressive Ratio 01 does not correlate with break point. **(E**.) Mean locomotion pre-lever during Long Access correlates with mean cocaine infusions during first 15 minutes of Long Access. **(F.)** Locomotion pre-lever during Progressive Ratio 02 does not correlate with break point. **(G.)** Locomotion pre-lever during Shock does not correlate with cocaine infusions during first 15 minutes of Shock. **(H.)** Active lever entrances per meter pre-lever across experiment. \**p*<.05, \*\**p*<.01, \*\*\**p*<.001 vs ShA01. **(I.)** Comparison of abstinence levels within and between session types. Within session type: \*\*\**p*<.001 vs 18 hours; between session type and abstinence: ###*p*<.001 vs 18, 66, and 90 hours Short Access; between session type: $$$*p*<.001 vs Short Access. (J.) Mean active lever entrances per meter pre-lever during Short Access correlates with mean cocaine infusions during first 15 minutes of Short Access. **(K.)** Active lever entrances per meter pre-lever during Progressive Ratio 01 correlates with break point. **(L**.) Mean active lever entrances per meter pre-lever during Long Access correlates with mean cocaine infusions during first 15 minutes of Long Access. **(M.)** Active lever entrances per meter pre-lever during Progressive Ratio 02 does not correlate with break point. **(N.)** Active lever entrances per meter pre-lever during Shock correlates with cocaine infusions during first 15 minutes of Shock. Plots A, B, H, and I represent mean and SEM. Correlation plots E and L are by Pearson, while remaining correlations are by Spearman Rank.

To assess the influence of session type and abstinence on locomotion pre-lever, a Type III Wald test was conducted following a 0.28 lambda Box-Cox transformation.

Significant main effects of session type (*χ*²(1)=11.93, *p*=.002), abstinence (*χ*²(2)=14, *p*=.002), and their interaction (*χ*²(2)=27.57, *p*<.001) were found. Within Short Access, post hoc analyses revealed locomotion pre-lever at 90 hours was significantly greater than 18 hours. Within Long Access, 66 and 90 hours was significantly greater than 18 hours. Interactively, locomotion pre-lever significantly decreased at 18 hours from Short to Long Access. At 66 hours, there was a significant increase from Short to Long Access. Overall, locomotion pre-lever was significantly greater in Long Access than in Short Access (**Fig. 3B**). These results show that locomotion pre-lever increases with drug history and hours of abstinence.

To assess locomotion pre-lever predictability of cocaine intake in the first 15 minutes of the session and break point, correlation analyses were performed. Between locomotion pre-lever and cocaine infusions during the first 15 minutes, a positive correlation was found during Short Access (*r_s_*(4222)=.5, *p*=.002) (**Fig. 3C**), and Long Access (*r*(4.29)=.59, *p*<.001, 95% CI [0.33, 0.77]) (**Fig. 3E**), but not during Shock (*r-_s_*(6952)=.18, *p*=.298) (**Fig. 3G**). Between locomotion pre-lever and breaking point, no correlations were found in both Progressive Ratio 01 (*r_s_*(7497)=.04, *p*=.84) or Progressive Ratio 02 (*r_s_*(6689)=.21, *p*=.219) (**Fig. 3D & 3F**). These results indicate that locomotion pre-lever is predictive of cocaine intake during the first 15 minutes in Short and Long Access, but is not predictive of breaking point.

#### 3.3.2. Active lever entrances per meter pre-lever escalate from Short to Long Access and predict cocaine intake during the first 15 minutes of self-administration

Initial analyses indicated that nose entrances into the active lever area may be predictive of intake. However, there was large individual variability in locomotion. To account for animals with higher general activity having a greater probability of entering the active lever area by chance, active lever area entrances were normalized by locomotion. To determine if active lever entrances per meter increased over the experiment, a Type III Wald test was performed. A significant main effect of session was found (*χ*²(23)=234.44, *p*<.001). Post hoc comparisons with Short Access 01 revealed significant increases in all sessions from Short Access 07 to Long Access 14, excluding Long Access 01, 03, and 07 (**Fig. 3H**). These results confirm that active lever entrances per meter increased over the experiment.

To assess the effects of session type and abstinence on active lever entrances per meter, a Type III Wald test was performed. No effect of abstinence was found (*χ*²(2)=1.7, *p*=.89), but a significant main effect of session type (*χ*²(1)=13.66, *p*<.001), and their interaction (*χ*²(2)=17.47, *p*<.001). Post hoc analyses revealed no significant differences between abstinence levels within Short Access. Within Long Access, significant increases were found at 66 and 90 hours in comparison to 18 hours.

Interactively, at every level of abstinence there was found to be a significant increase from Short to Long Access (**Fig. 3I**). These results confirm that active lever entrances per meter increase with drug experience, particularly after longer periods of abstinence.

To assess active lever entrances per meter predictability of cocaine intake in the first 15 minutes of the session and break point, correlation analyses were performed.

Between active lever entrances per meter and cocaine infusions during the first 15 minutes, a significant positive correlation was found during Short Access (*r_s_*(3002)=.64, *p*<.001) (**Fig. 3J**), Long Access (*r*(4.65)=.62, *p*<.001, 95% CI [0.37, 0.79]) (**Fig. 3L**), and Shock (*r_s_*(5199)=.38, *p*=.019) (**Fig. 3N**). Between active lever entrances per meter and break point, a significant positive correlation was found during Progressive Ratio 01 (*r-_s_*(5173)=.33, *p*=.046) (**Fig. 3K**), but not Progressive Ratio 02 (*r_s_*(7985)=.05, *p*=.753) (**Fig. 3M**). These results show that active lever entrances per meter pre-lever are predictive of both cocaine intake within the first 15 minutes and motivation measured as break point following Short Access, but not after Long Access.

### 3.4. Lever Entrances During Noncontingent Sessions

#### 3.4.1. Lever entrances per meter show a preference for the active lever after a history of self-administration

To determine if seeking behavior was expressed during Noncontingent sessions, entrances per meter into the active and inactive lever areas were analyzed. Considering the levers never extend during Noncontingent sessions they were treated as pre-lever sessions. Examination of raw data showed low variation within Noncontingent sessions and therefore all three sessions (e.g. Baseline, Saline, and Drug) were grouped for analyses. Noncontingent 01 was then treated as the Baseline, and Noncontingent 02 and 03 active and inactive lever entrances per meter were normalized by the group mean of Noncontingent 01 to account for high individual variation. Following a 0.23 lambda Yeo-Johnson power transformation, a Type III Wald test indicated a significant main effect of session (*χ*²(1)=12.46, *p*=.002), but not a significant main effect of levers (*χ*²(1)=3.12, *p=*.162) or their interaction (*χ*²(1)=3.49, *p*=.162). Post hoc analyses revealed Noncontingent 03 to be significantly greater than Noncontingent 02, prompting a paired Student’s t-test between the active and inactive levers in Noncontingent 03 where active was found to be greater than inactive. Direct comparisons to Noncontingent 01 found that the inactive lever entrances per meter in Noncontingent 02 were significantly less than baseline, while the active lever entrances per meter in Noncontingent 03 were significantly greater than baseline. (**Fig. 4A**). These results indicate that by Noncontingent 03 rats demonstrate preferential seeking behavior toward the drug-associated active lever area.

**Figure 4.**
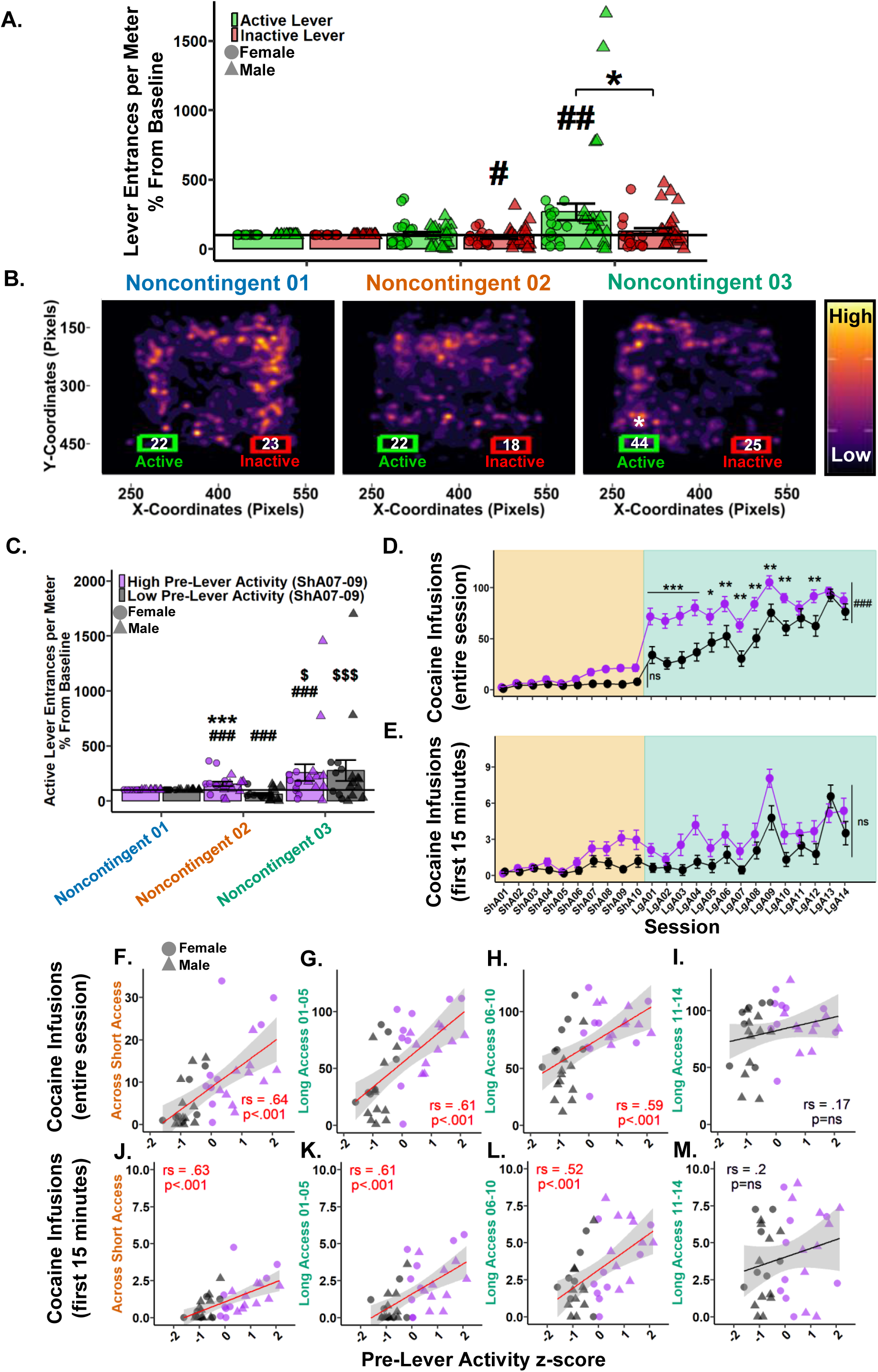
Pre-lever activity during Short Access 07-09 predicts cocaine intake in Short and Long Access and seeking behavior during Noncontingent sessions. **(A.)** Active (green) and inactive (red) lever entrances per meter as percentage difference from Baseline (Noncontingent 01). *#p*<.05, *##p*<.01 vs Baseline; \**p*<.05 vs inactive lever. **(B.)** Nose (*x, y)* pixel coordinate heatmaps of all rats within each Noncontingent session in relation to active (green) and inactive (red) Levers. Each plot represents ∼109 sessions overlain revealing high and low activity zones. Numbers within lever bounding boxes are the mean sum of active and inactive lever entrances for all rats within each Noncontingent session (Noncontingent 01 active: *M*=22.2, *SD*=13.9; inactive: *M*=23.4, *SD*=12.3; Noncontingent 02 active: *M*=21.8, *SD*=16.1; inactive: *M*=18.4, *SD*=16.5; Noncontingent 03 active: *M*=43.5, *SD*=42.1; inactive: *M*=24.7, *SD*=22.1). \**p<*.05 vs inactive lever. **(C.)** Active lever entrances per meter percent difference from Baseline (Noncontingent 01) grouped by pre-lever activity z-score during Short Access 07-09. *###p*<.001 vs Baseline; Within pre-lever activity group: $*p*<.05, $$$*p*<.001 vs Noncontingent 02; Between pre-lever activity groups: \*\*\**p*<.001 vs Noncontingent 02 Low group. **(D-E.)** Cocaine infusions over the entire session (D.) and within the first 15 minutes (E.) grouped by Short Access 07-09 pre-lever activity z-score. \**p*<.05, \*\**p*<.01, \*\*\**p*<.001, ###*p*<.001 vs Low. **(F-I.)** Cocaine infusions over the entire session correlated using Spearman Rank with Short Access 07-09 pre-lever activity z-score in Short Access (F.), Long Access 01-05 (G.), Long Access 06-10 (H.), but not during Long Access 11-14 (I.). **(J-M.)** Cocaine infusions within the first 15 minutes correlated using Spearman Rank with Short Access 07-09 pre-lever activity z-score in Short Access (J.), Long Access 01-05 (K.), Long Access 06-10 (L.), but not during Long Access 11-14 (M.). Plots represent means and SEM.

To visualize the development of interest in the active lever, density heat maps of each rat’s nose node coordinates (*x*, *y*) for all sessions within each Noncontingent session were overlain. Bounding boxes corresponding to the active and inactive lever areas were plotted with the rounded mean lever entrances. Student’s t-tests between active and inactive lever entrances within Noncontingent sessions revealed only Noncontingent 03 following a 0.23 lambda Yeo-Johnson power transformation achieved significance, with active greater than inactive (**Fig. 4B**). These results highlight the development of interest in the active lever after a history or self-administration as the high activity regions migrate to the active lever side by Noncontingent 03.

#### 3.4.2. Pre-lever activity during Short Access sessions 07-09 predicts active lever entrances per meter during Noncontingent sessions

To determine if the interest in the active lever was developed during self-administration, rats were grouped as “High” (n=19) or “Low” (n=18) by a median-split z-score of their pre-lever activity measures (locomotion and active lever entrances per meter) during Short Access 07-09. These sessions were chosen as they were the first to show significance compared to Short Access 01 and are the last Short Access sessions before Noncontingent 02. First, the average of each variable was computed across the three sessions before being standardized within the respective variables. The two z-scores were then averaged, where higher scores equate to greater pre-session activity. Following a median z-score split, those greater than or equal to the median were deemed “High”, and those less than were deemed “Low”.

Following a 0.2 lambda Yeo-Johnson power transformation, a Type III Wald test indicated no main effect of session (*χ*²(1)=4.81, *p*=.059), but a significant effect of pre-lever activity (*χ*²(1)=29.53, *p*<.001) and their interaction (*χ*²(1)=9.39, *p*=.006). Post hoc analyses revealed the High pre-lever activity group had significantly greater active lever entrances per meter in both Noncontingent 02 and 03 than baseline and versus the Low pre-lever activity group within Noncontingent 02. For the Low pre-lever activity group, Noncontingent 02 was significantly less than both baseline and Noncontingent 03.

There was no difference between groups in Noncontingent 03 (**Fig. 4C**). These results confirm that rats with high pre-lever activity during Short Access 07-09 led to significant seeking in Noncontingent 02 that persisted in Noncontingent 03. Furthermore, the Low pre-lever activity group developed an interest during Long Access before Noncontingent 03.

#### 3.4.3. Pre-lever activity during Short Access sessions 07-09 predicts cocaine self-administration in Short and Long Access

To determine if pre-lever activity during Short Access sessions 07-09 predicts cocaine intake over the entire session in Short Access, a Type III Wald test was performed. A significant main effect was found in session (*χ*²(9)=125.52, *p*<.001), but not a significant effect of pre-lever activity (*χ*²(1)=3.51, *p*=.169) or their interaction (*χ*²(9)=16.9, *p*=.169) (**Fig. 4D**). These results show that cocaine intake did not differ between groups in Short Access.

To determine if pre-lever activity during Short Access sessions 07-09 predicts cocaine intake over the entire session in Long Access, a Type III Wald test was performed. Significant main effects were found in session (*χ*²(13)=62.03, *p*<.001), pre-lever activity (*χ*²(1)=13.49, *p*<.001) and their interaction (*χ*²(13)=28.79, *p*=.015). Post hoc analyses revealed the High pre-lever activity group to have significantly greater cocaine infusions over the entire session in all Long Access sessions except Long Access 11, 13, and 14 (**Fig. 4D**). These results confirm that pre-lever activity during Short Access 07-09 was initially predictive of cocaine intake during Long Access, but that groups eventually converged.

To determine if pre-lever activity during Short Access sessions 07-09 predicts cocaine intake within the first 15 minutes of the session, a Type III Wald test was performed. A significant main effect was found in session (*χ*²(23)=208.9, *p*<.001), but not in pre-lever activity (*χ*²(1)=0.85, *p*=.742) or their interaction (*χ*²(23)=35.32, *p*=.134) (**Fig 4E**). These results suggest that pre-lever activity during Short Access 07-09 is not predictive of loading behavior between groups.

Considering the incongruency of the significant association between pre-lever variables and cocaine intake variables (**Fig. 3**), and the previous nonsignificant results between pre-lever activity groups and cocaine infusions in the first 15 minutes (**Fig. 4D**), correlations were performed between cocaine intake variables and pre-lever activity z-scores in Short Access and three separate phases of Long Access. For cocaine infusions during the entire session and pre-lever activity, significant positive correlations were found in Short Access (*r_s_*(2960)=.65, *p*<.001; **Fig. 4F**), Long Access 01-05 (*r-_s_*(3284)=.61, *p*<.001; **Fig. 4G**), Long Access 06-10 (*r_s_*(3422)=.59, *p*<.001; **Fig. 4H**), but not during Long Access 11-14 (*r_s_*(7013)=.17, *p*=.318; **Fig. 4I**). For cocaine infusions during the first 15 minutes and pre-lever activity, significant positive correlations were similarly found in Short Access (*r_s_*(3094)=.63, *p*<.001; **Fig. 4J**), Long Access 01-05 (*r_s_* (3301)=.61, *p*<.001; **Fig. 4K**), Long Access 06-10 (*r_s_*(4023)=.52, *p*<.001; **Fig. 4L**), but not during Long Access 11-14 (*r_s_*(6735)=.2, *p*=.231; **Fig. 4M**). These results confirm that rats who showed markedly increased pre-lever activity during Short Access sessions 07-09 were more likely to consume more cocaine during the first 15 minutes and over the entire session in both Short and Long Access, but converged by the end of Long Access. Furthermore, these results support those shown in **Fig. 4C**, where no significant difference was found in active lever entrances per meter in Noncontingent 03, which occurs after Long Access 14.

## 4. Discussion

Here, we demonstrate that cocaine infusions over the full session and within the first 15 minutes escalated during Short and Long Access in heterogeneous stock rats. Locomotion increased after a noncontingent cocaine infusion, and population-level analysis showed no signs of psychomotor sensitization or tolerance following Short Access (Noncontingent 02) or Long Access (Noncontingent 03). Individual analyses revealed strong divergence in psychomotor sensitization or tolerance expression, though neither influenced cocaine self-administration. Pre-lever locomotion and active lever entrances per meter escalated from Short to Long Access, increased after instances of abstinence, and predicted early session intake. Rats with high pre-lever activity during Short Access sessions 07-09 escalated active lever entrances across Noncontingent sessions, and earned more infusions during Short Access and first two weeks of Long Access than low pre-lever activity rats, but these differences disappeared by Long Access 13 as low-activity rats escalated to similar levels. Together, the data dissociate psychomotor sensitization and tolerance from cocaine intake and identify pre-lever activity as a robust predictor of cocaine intake and escalation.

### 4.1. Cocaine Intake

Rats increased total cocaine intake as early as Short Access 02 and within the first 15 minutes as early as Short Access 07, while exhibiting significant escalation of intake after two weeks of Long Access. These findings align with our previous reports that heterogenous stock rats show increased cocaine intake during Short and Long Access [28], and the larger literature that Long Access induces escalation of intake [3,4,24,55].

### 4.2. Psychomotor Expression

Population-level analyses revealed locomotion increased following a noncontingent infusion of cocaine, but no differences between Drug sessions. Saline 02 was significantly greater than Baseline 02, suggesting an anticipatory conditioned stimulus effect as the last time they were handled as such they received a noncontingent infusion of cocaine (Drug 01). However, the effect is not seen following Long Access, suggesting that this conditioned responding is short-lasting and unlikely to drive escalation of cocaine self-administration. Individually, rats did show differences in the propensity to develop psychomotor sensitization or tolerance to noncontingent cocaine administration. One interpretation is that analyzing only ambulatory activity may misclassify some rats displaying stereotypic behavior, such as head bobbing, weaving, sniffing, or gnawing at high cocaine doses [8,56]. However, this is unlikely for several reasons. First, the dose used (1.5 mg/kg) is known to produce minimal stereotypic behavior and robust sensitization [6,30–32]. Second, the nose motion (a proxy for stereotypic behavior) was not found to increase in Tolerant rats (**Fig. S1**). Third, visual inspection of the videos show that the most tolerant rat (M2677) did not exhibit stereotypic behavior while the most sensitized rat (M2668) showed pronounced psychomotor sensitization. A possible explanation for the difference between sensitized vs. tolerant rats is that sensitized rats consumed less cocaine, while tolerant rats consumed more. However, intake analyses do not support this hypothesis: groups showed no differences in cocaine intake in Short or Long Access, or within the first 15 minutes. These findings challenge the idea that Short Access produces only sensitization and Long Access only tolerance to the effects of cocaine [1,3–6,18,24,57], and show that neither reliably predicts cocaine intake.

### 4.3. Pre-Lever Activity Measures

Pre-lever variables locomotion and active lever entrances per meter increased from Short to Long Access and intensified following two-to-three days of abstinence. Similarly, rats showed a significant increase in the active lever area entrances per meter in Noncontingent 03 in comparison to Noncontingent 01, suggesting sensitization of drug-seeking behavior. In contrast to measures of psychomotor sensitization or tolerance, both locomotion and active lever entrances per meter were highly predictive of intake. These findings align with the incentive salience literature, showing that repeated drug use leads to salience attribution to cues and environments that particularly manifests following a period of abstinence [5–6,15–16,19–20,30–32,46–47,58–65].

### 4.4. Individual Differences in Pre-Lever Activity Measures

To investigate the individual propensity to attribute salience to drug-associated cues, a median z-score of pre-lever activity was calculated. Short Access sessions 07–09 were selected as they occurred within the same week (no abstinence periods) and were the first sessions in which both pre-lever variables and infusions within the first 15 minutes reached significance (**Fig. 1C, 3A, & 3H**). Comparisons between High and Low pre-lever activity groups revealed that high-activity rats exhibited an early escalation of cocaine intake during Long Access, but ultimately both pre-lever activity groups reached a ceiling level of intake after two weeks when groups no longer differed.

One possible explanation is that Low pre-lever activity rats in Short Access were slower in developing incentive salience, and once expressed, escalated their intake to levels comparable to High pre-lever activity rats. Indeed, the significant difference in active lever entrances per meter between pre-lever activity groups in Noncontingent 02 and its absence in Noncontingent 03 supports this interpretation. These findings align with previous reports that individuals differ in their susceptibility to attribute incentive salience, which may serve as an early predictor of vulnerability [56,59,62–63].

Furthermore, they suggest that incentive salience is a catalyst of escalation, contrary to previous findings that incentive salience and escalation are dissociable [26].

Another interpretation is that escalating pre-lever activity during Long Access reflects the combined influence of allostatic load [1,3,55,57], incentive habit formation [66–68], and the incubation of craving [69–70]. Allostasis theory predicts that repeated use progressively recruits stress systems and negative affect, shifting the hedonic set point and motivating drug seeking to alleviate withdrawal-related distress. Consistent with this view, both animal and human studies show that acute withdrawal, withdrawal-conditioned cues, negative affect, and psychosocial stress increase craving, invigorate drug seeking, and amplify cue-evoked neural activity [1,3,55,57,71–75]. The incentive habit framework suggests that repeated drug-cue pairing transforms early cue reactivity into motivationally charged habitual responses, which may lead to automatic approach toward drug-associated cues even before drug availability [66–68]. Finally, incubation of craving provides an additional, complementary mechanism: periods of abstinence without obvious negative emotional states enhance the motivational impact of drug cues over time, strengthening cue-triggered approach behavior [69–70]. Follow up studies will be needed to dissect the role of each of these processes.

The high pre-lever activity phenotype we observed shares conceptual parallels with the sign-tracker/goal-tracker framework [59]. Sign-trackers preferentially interact with a discrete cue (the lever CS), while goal-trackers orient to the site of reward delivery. Sign-trackers have been shown to initially self-administer more cocaine [63] and choose cocaine over food [76]. However, important differences exist: sign-tracking is defined in a purely Pavlovian context, with lever pressing having no instrumental consequence, whereas our measure reflects operant-conditioned lever-directed exploration. Notably, recent work has shown that sign-/goal-tracker status does not reliably predict overall cocaine self-administration or escalation, although sign-trackers may show greater resistance to punishment [77]. Thus, our results extend the sign-/goal-tracker framework into an operant cocaine paradigm, highlighting that cue-bias and responsivity can shape early intake and drive escalation of intake with prolonged access even in initially resistant animals.

### 4.5. Limitations

The present study has a few limitations. One is the lack of a control group without periods of abstinence. A control group participating in successive sessions could further confirm the influence of abstinence on incentive salience incubation. However, noncontingent sessions 02 and 03 did not follow a period of abstinence, and expressions of incentive salience and both psychomotor sensitization and tolerance were still observed. Another limitation is the use of pose-estimation over manual scoring of behavior could result in inaccuracies. However, pose-estimation with SLEAP is remarkably accurate [37,78], and the instance rendered sample videos display precise tracking of the catheter and nose nodes (see supplementary videos).

### 4.6. Summary

In summary, this report demonstrates that individual differences in psychomotor sensitization or tolerance had no influence on cocaine intake. In contrast, incentive salience during Short Access acts as a catalyst and early predictor of escalation in cocaine self-administration during the first two weeks of Long Access, but does not predict established escalation of cocaine intake after three weeks of Long Access.

Moreover, individuals initially resistant to incentive salience can, with sufficient exposure, become sensitized and escalate cocaine use to the same level as more susceptible individuals. Expression of incentive salience sensitization as reflected in pre-lever activity measures and their subsequent influence on cocaine intake supports the idea that incentive salience and psychomotor sensitization are mediated by distinct mechanisms [6–7,21,31]. The development of the incentive salience profiles is accelerated by periods of abstinence (weekend), which allows for incubation of craving, a progressive increase of craving during early days of abstinence that can remain elevated following prolonged abstinence [44,69–70]. The measure of approach toward the active lever area using video recording before the start of the self-administration session may represent a novel behavioral marker of addiction vulnerability that can precisely track the motivation for the drug with high accuracy without using any operant measures, the drug itself, or any other training or conditioning that could interfere with the drug self-administration. Such a measure may also be useful for medication development to allow dissociation between drug-dependent and drug-independent effects of experimental medication on drug seeking and taking.

## Supporting information

Mixed Model Fitting Process and Sex Effects

Final Mixed Models for Analyses

Sensitized M2668 and Tolerant M2677 Rendered Videos

## Data Availability Statement

The datasets generated during and/or analyzed during the current study are available from the corresponding author on reasonable request.

## Acknowledgements

This research was done using services provided by the OSG Consortium [38–41] which is supported by the National Science Foundation awards #2030508 and #1836650.

## Author Contributions

Jarryd Ramborger: Manuscript drafting and revising, data collection, experiment execution, experimental design, data analysis and interpretation

Joseph Mosquera: Data collection, experiment execution

Molly Brenna: Data collection, experiment execution

Benjamin Sichel: Data collection, experiment execution

Dyar Othman: Data collection, experiment execution

Sonja Plasil, PhD: Data collection, experiment execution

Elizabeth Sneddon, PhD: Data collection, experiment execution

Selene Zahedi: Data collection, experiment execution

Alex Morgan: Data collection, experiment execution

Supakorn Chonwattanagul: Data collection, experiment execution

Kathleen Bai: Data collection, experiment execution

Lindsey China: Data collection, experiment execution

Tu La: Data collection, experiment execution

Lisa Maturin: Data collection

Lieselot L. G. Carrette, PhD: Experimental design, manuscript drafting and revising

Olivier George, PhD: Manuscript drafting and revising, experimental design, data analysis and interpretation, funding and resources, final approval

