## Supplementary material for "Incentive Salience, not Psychomotor Sensitization or Tolerance, Drives Escalation of Cocaine Self-Administration in Heterogeneous Stock Rats": Mixed Model Fitting Process and Sex Effects

### Introduction

The following document provides a detailed account of the model fitting and choice process for the linear and generalized mixed models utilized in the repeated measure ANOVA statistical analyses, a test of stereotypies to accompany **Fig. 2C**, and analyses with sex as a factor. First is the step-by-step process of testing and comparing models with different model fits. Next are the assumption test visualizations provided by the simulateResiduals() function from the RStudio DHARMa package (Hartig, 2024) for each final mixed model, including the analyses of sex and stereotypies. The function had the parameters “refit = FALSE” and “n = 500” for residual simulations. The default number of simulations is 250, and 500 was chosen for computation time and greater precision (Hartig, 2024). Each plot is identified alphabetically for how they appear in the manuscript, and each model’s statistics can be found in the excel spreadsheet table. Next are the results and figure for the stereotypies analysis. And lastly are the analyses with sex included as a factor.

### 1. Model Testing and Comparison of Fit Process

To best explain the process of testing and comparing different linear and generalized linear mixed model combinations, an example is used to fit the steps in context. The chosen example is the first statistical analysis in the manuscript and supplementary excel spreadsheet, i.e., answering the question: “Do cocaine infusions in the entire session differ between sessions in Short Access”? Therefore, the response variable is Cocaine Infusions, the fixed effect is Session, and given this is a repeated measure analysis, the random effect is each individual Rat. The formula, as it would appear in RStudio, would then be:

*response variable ~ fixed effects + random effects*

or in this example,

*Cocaine Infusions ~ Session + (1|Rat)*

This formula is then used to fit a model of the data using the linear mixed effects model function lmer() (Bates et al., 2015; Kuznetsova, Brockhoff, & Christensen, 2017) or a generalized linear mixed effects model with glmmTMB() (Brooks et al., 2017). All models were fit with maximum likelihood to allow valid statistical comparisons of fixed effects structures and to support post hoc testing as recommended (Bates et al., 2015; Luke, 2017; Meteyard & Davies, 2020). The process is as follows:

1. The first step was to determine the optimal lambda for Box-Cox of Yeo-Johnson power transformations to achieve normality for a linear regression model. Box-Cox are for response variables that do not contain zero or negatives, and Yeo-Johnson for those that do. For this example, a Yeo-Johnson was performed.
2. Two linear mixed models using lmer() were then produced: one without power transformation, and one with power transformation.
3. The two linear mixed models were then compared with a likelihood ratio test.

4. The two linear mixed models were then visualized with simulateResiduals() in the order of the likelihood ratio test results, i.e., worst to best, and their results documented. These steps elucidate the modeling done by linear methods.
5. Next, a number of generalized linear mixed models are fit using glmmTMB. Unlike linear mixed models, generalized mixed models allow for specification of both the distribution family (e.g., gaussian, logarithmic, tweedie, etc.) and the dispersion structure, enabling more flexible modeling of data with non-normal distributions or heterogeneous variance. For example, some distribution families require nonnegative and whole numbers (negative binomial distributions), while others do not have such constraints (gaussian). For this example, six separate base models were fit to the data: without transformation, with a Yeo-Johnson transformation, with a tweedie distribution, and all three available negative binomial distributions.
6. Similarly to the linear mixed models, the generalized mixed models were then compared using a likelihood ratio test.
7. The generalized mixed models were then visualized with simulateResiduals() in the order of the likelihood ratio test results from worst to best. Steps 5-7 reveals the best family distribution to be used.
8. The best-fitting model from the previous step is then used as a baseline to evaluate alternative dispersion structures within the generalized linear mixed model. Whereas linear mixed models estimate only the conditional mean of the outcome variable (e.g., $Y \sim X$), generalized linear mixed models permit simultaneous modeling of both the mean and the variance. By default, variance

is assumed to follow the form dictated by the specified distributional family. However, the inclusion of dispersion parameters enables the modeling of heteroskedasticity by allowing fixed effects (e.g., session, session type, etc.) to account for variation in the residual variance. This step evaluates whether explicitly modeling dispersion improves overall model fit and residual behavior. In the case of this example, a few of the dispersion parameters fit were:

*Cocaine Infusions ~ Session + (1|Rat)*

*Dispersion = Session*

*Dispersion = Session + (1|Rat)*

*Dispersion = Abstinence*

*Dispersion = Abstinence + (1|Rat)*

These specifications allow the variance to vary as a function of Session, as a function of both Session and individual Rats, by each level of Abstinence, or Abstinence and each individual Rat.

9. Each generalized mixed model from step 8 was then compared using a likelihood ratio test.
10. Each model from step 8 was then visualized with simulateResiduals() in the order of the likelihood ratio test results. It is in this step that the best linear mixed model simulateResiduals() results from above in step 4 is compared with the generalized mixed model combinations.
11. The model in Step 10 that yielded the most favorable simulateResiduals() diagnostics was selected as the final model. Primary emphasis was placed on

the Kolmogorov–Smirnov test to assess residual uniformity and overall model fit to the data. However, in instances where warranted, a balanced interpretation of all diagnostic outputs—which include dispersion, outlier detection, and observed versus predicted relationships—was preferred, as the Kolmogorov–Smirnov test is known to be sensitive to large sample sizes, which may yield statistically significant deviations despite an acceptable practical fit (Hartig, 2024; Razali & Wah, 2011).

### 2. DHARMa simulateResiduals() Assumption Check Results

Below are the plotted outputs generated by simulateResiduals() for the final model used in each of the repeated measure statistical analyses found in the manuscript. Red text indicates a failure of an assumption test. The plots are alphabetical in the order in which they appear in the manuscript and in the accompanying supplementary excel spreadsheet. The excel spreadsheet contains a table with the final formula of the models as well as the statistical results generated by summary(). Results of models not selected are available from the corresponding author on reasonable request.

**A. Cocaine Infusions in Short Access**

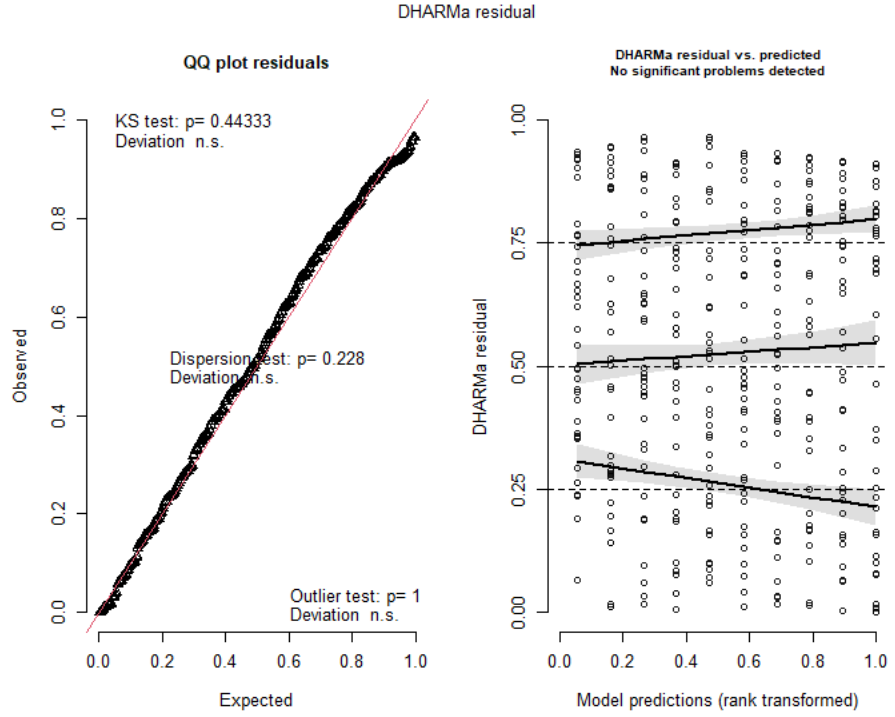


**B. Cocaine Infusions in Long Access**

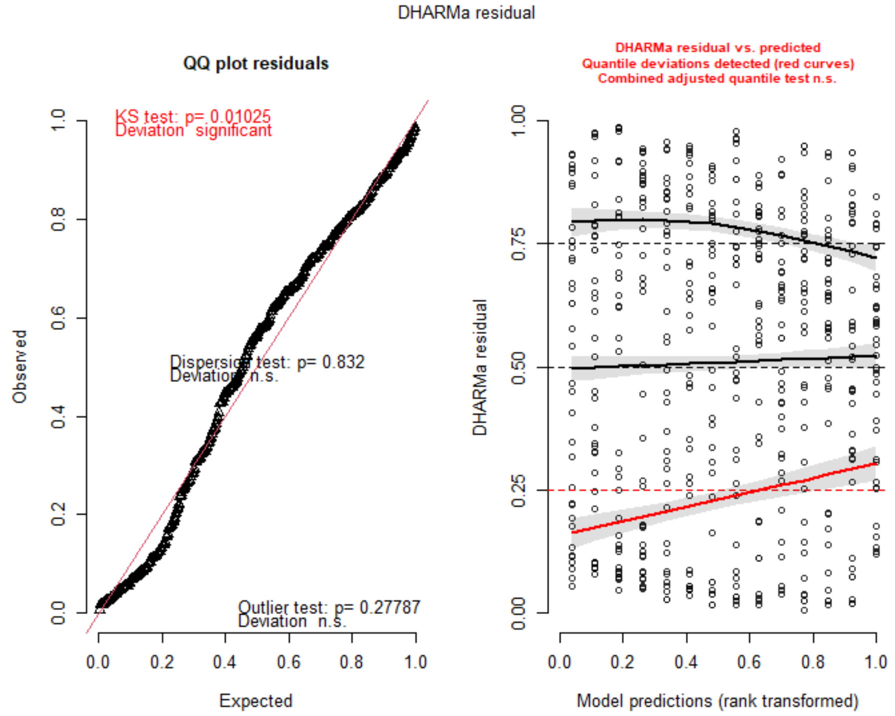

### C. Cocaine Infusions First Fifteen Minutes

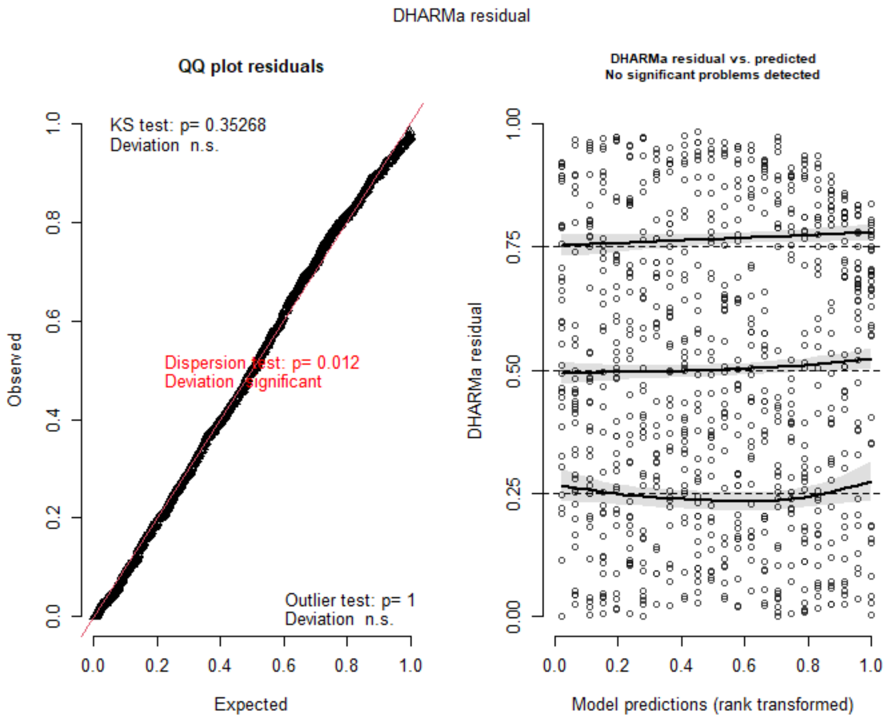

### D. Locomotion Raw in Noncontingent Sessions

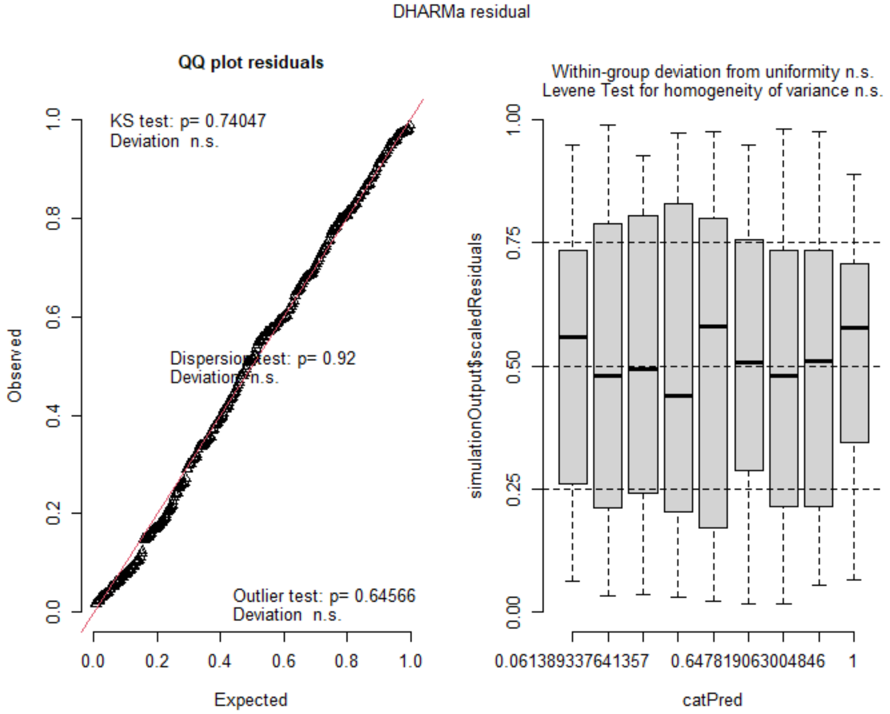


### E. Locomotion Percent Difference From Baseline in Noncontingent Sessions

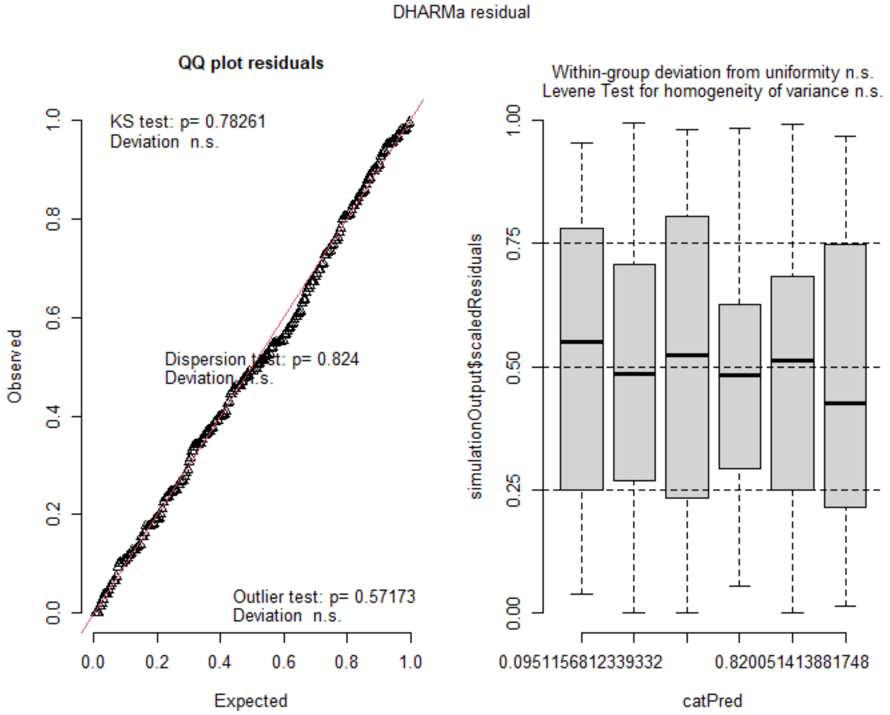

### F. Locomotion Percent Difference From Drug 01 by Behavioral Expression

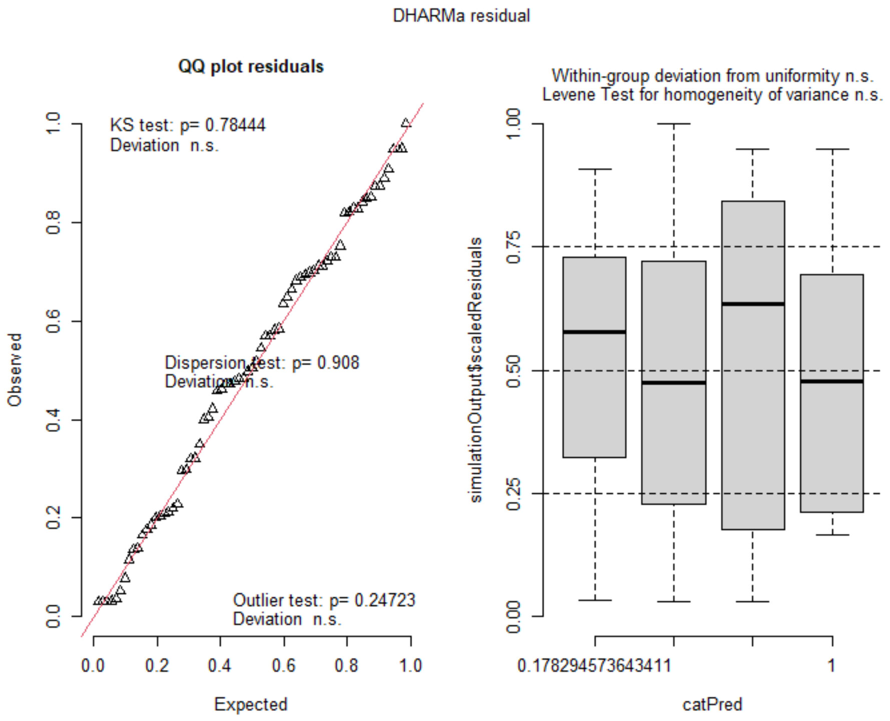


### G. Cocaine Infusions in Short Access by Behavioral Expression

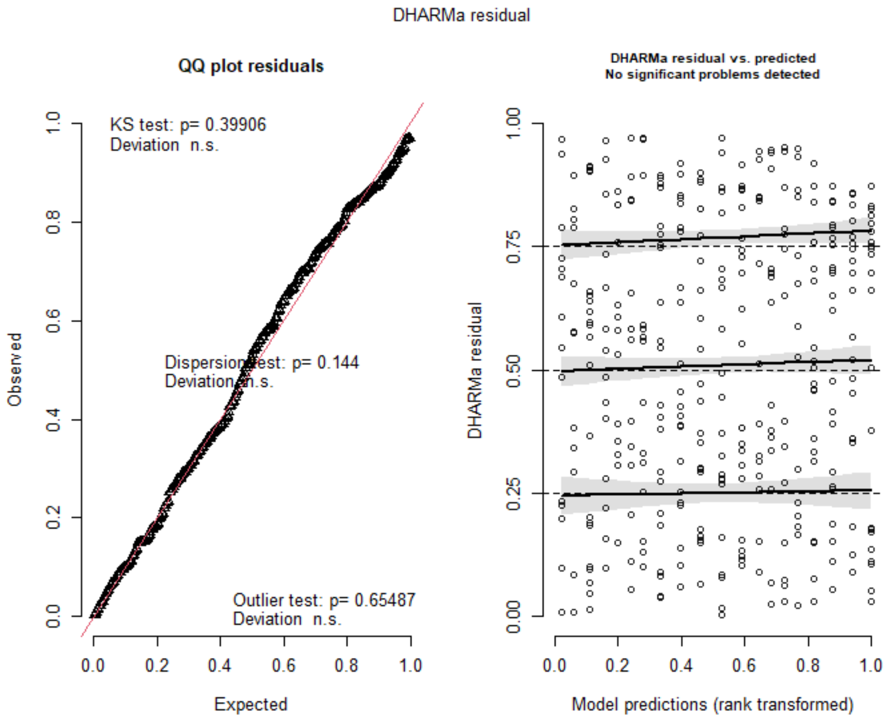

### H. Cocaine Infusions in Long Access by Behavioral Expression

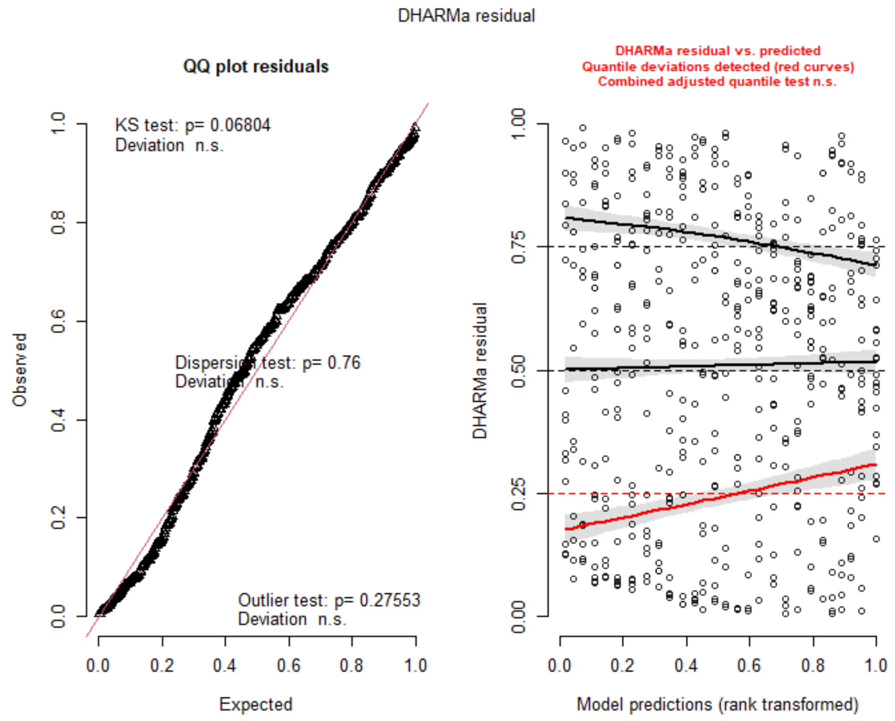


### I. Cocaine Infusions First 15 Minutes by Behavioral Expression

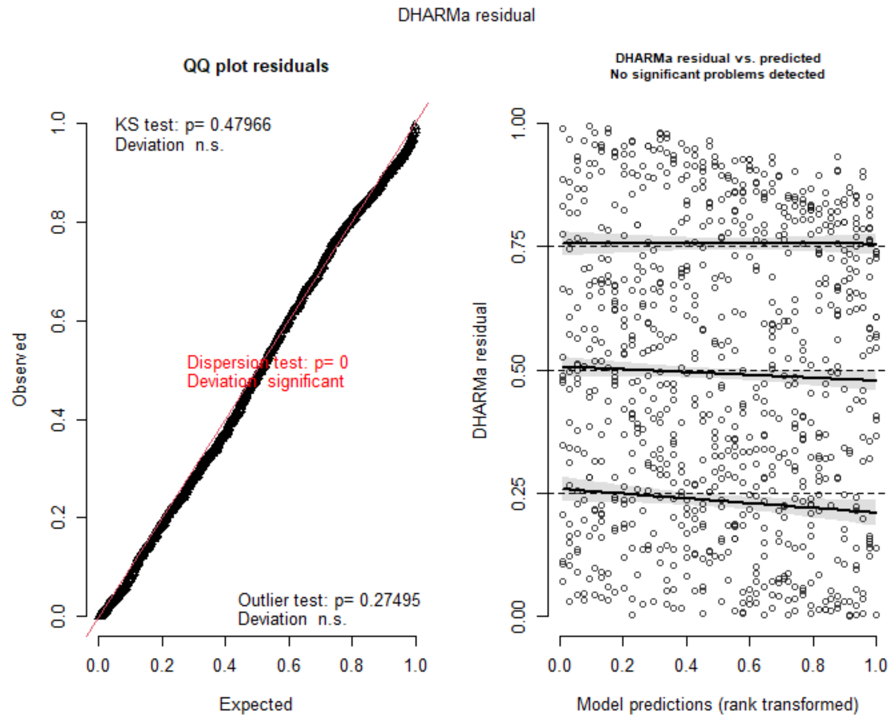

### J. Locomotion Pre-Lever

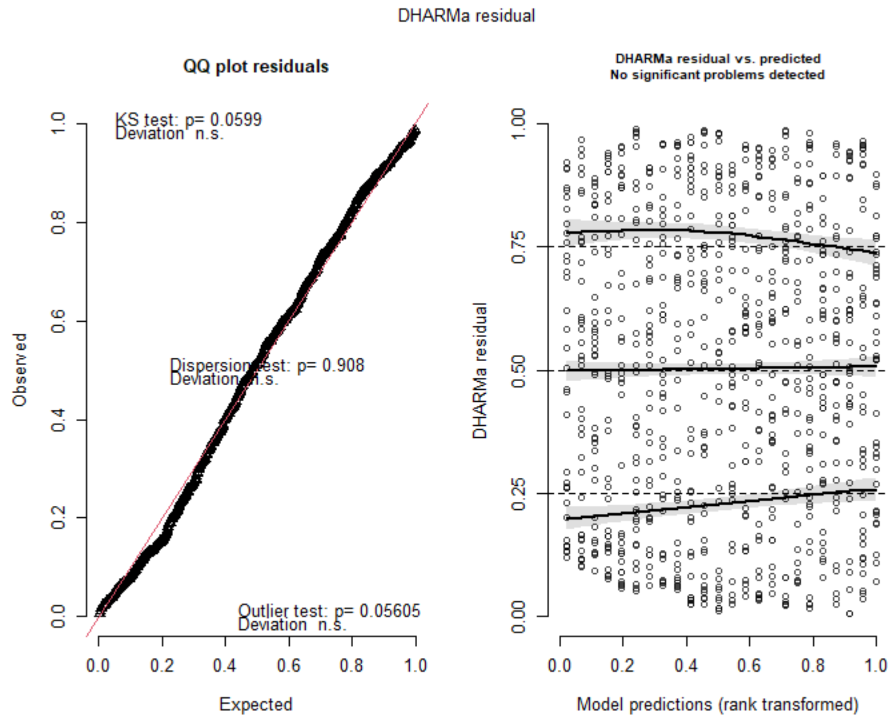


### K. Locomotion Pre-Lever by Session Type and Abstinence

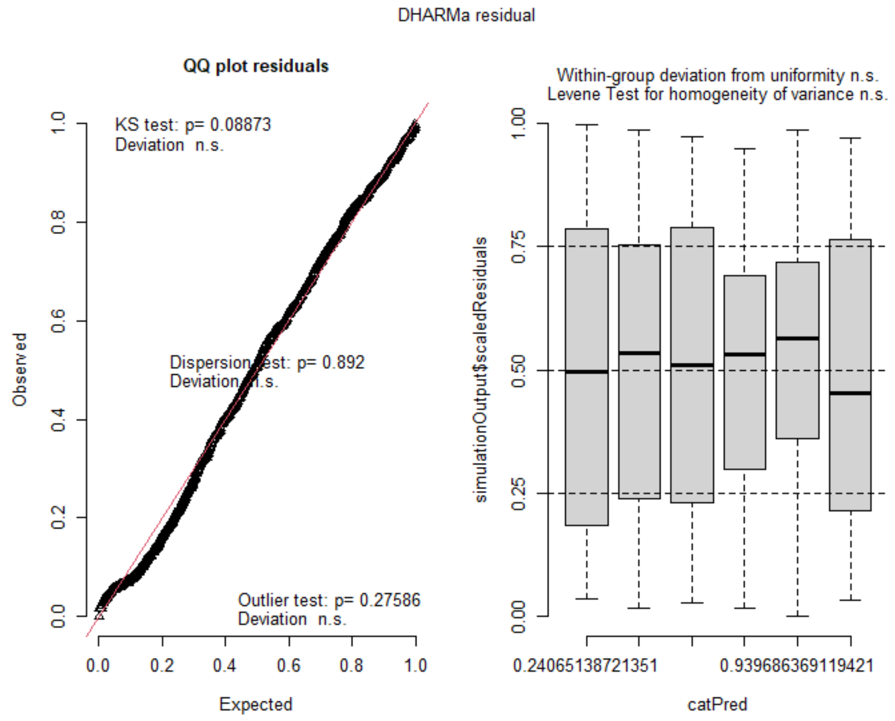

### L. Active Lever Entrances per Meter Pre-Lever

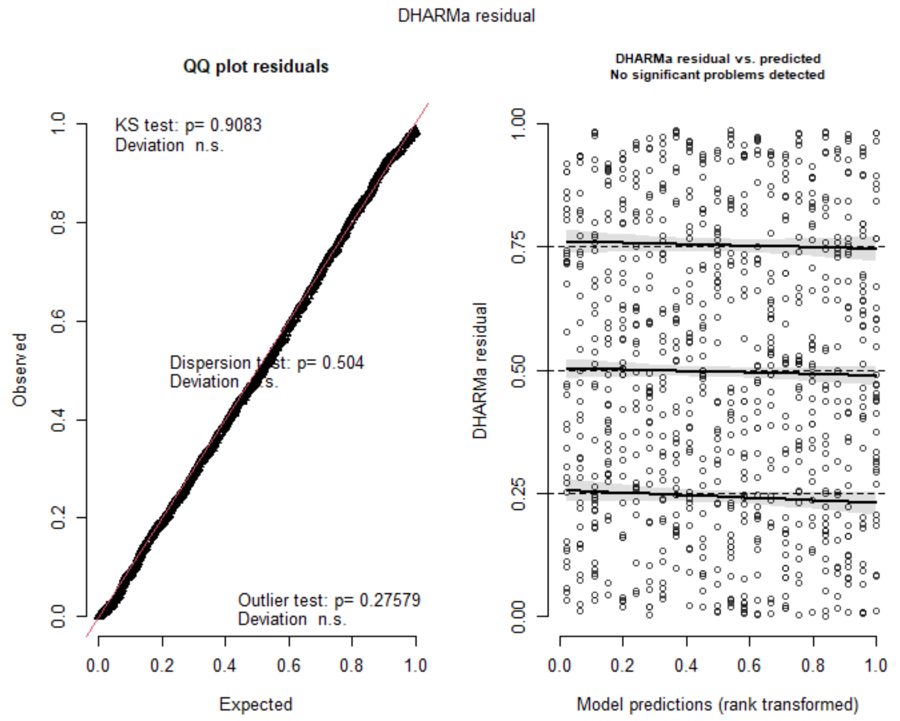


### M. Active Lever Entrances per Meter by Session Type and Abstinence

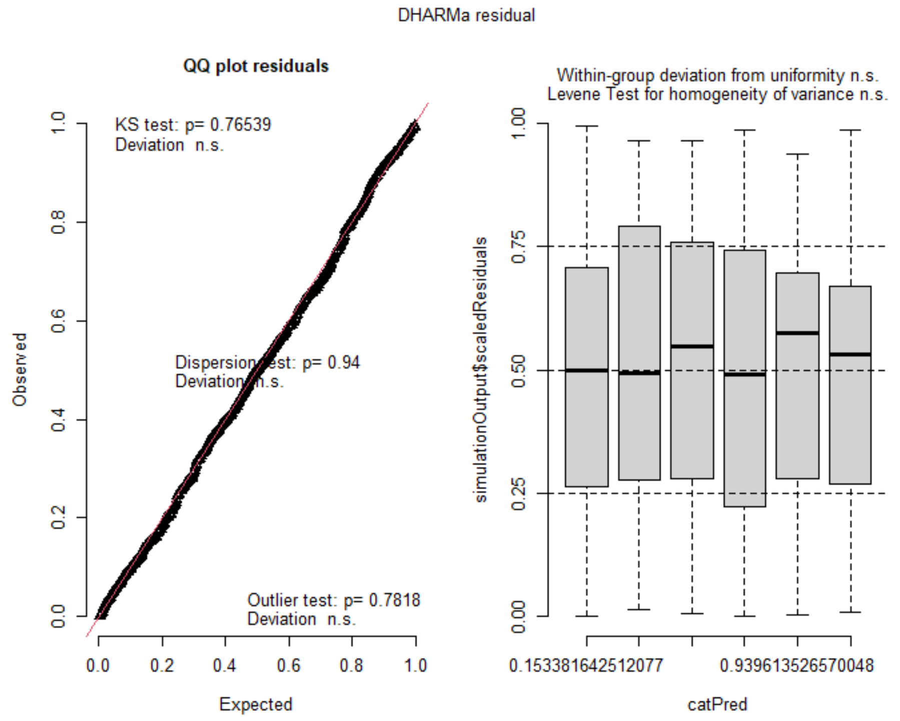

### N. Entrances per Meter Percent Difference From Noncontingent 01

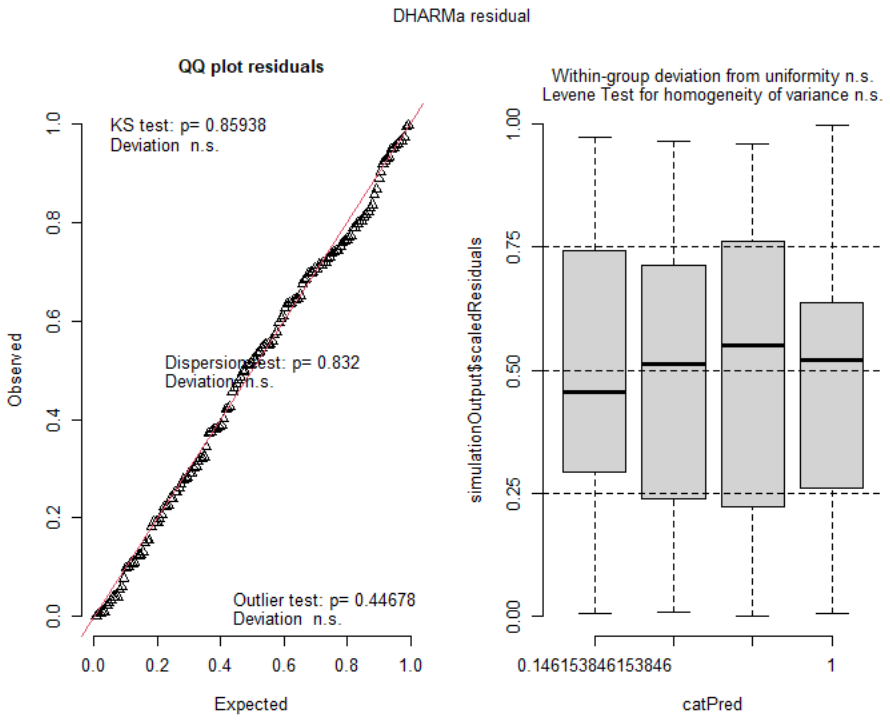


### O. Active Lever Entrances per Meter Percent Difference From Noncontingent 01 by Pre-Lever Activity

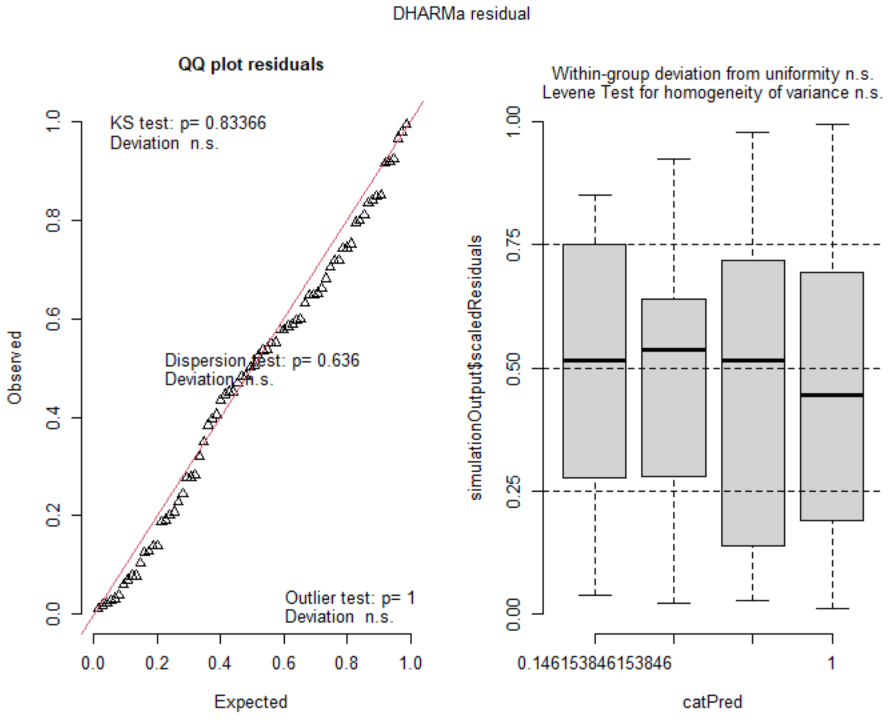

### P.  Cocaine Infusions in Short Access by Pre-Lever Activity

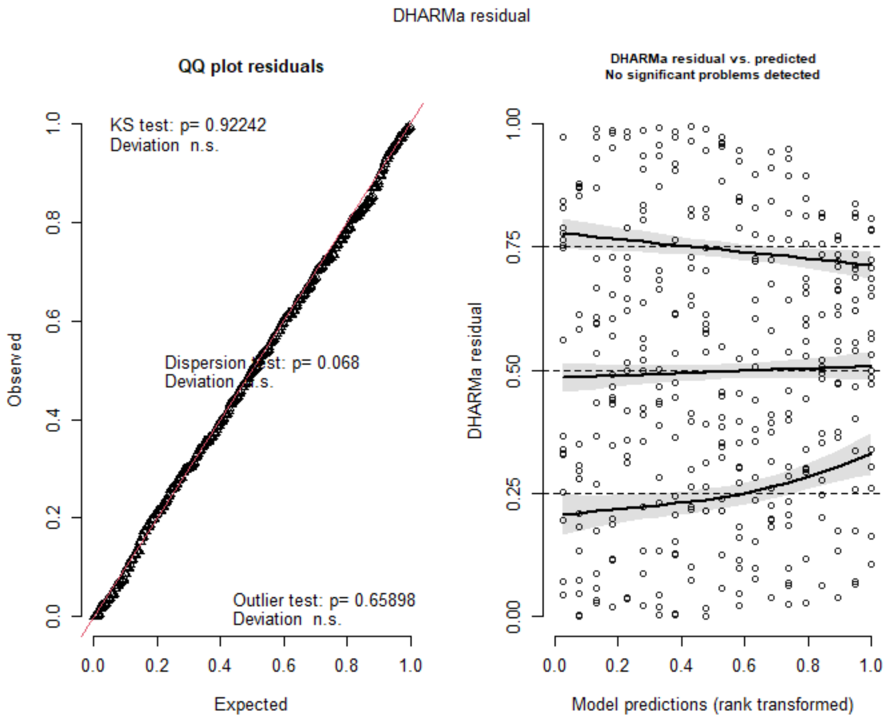


### Q.  Cocaine Infusions in Long Access by Pre-Lever Activity

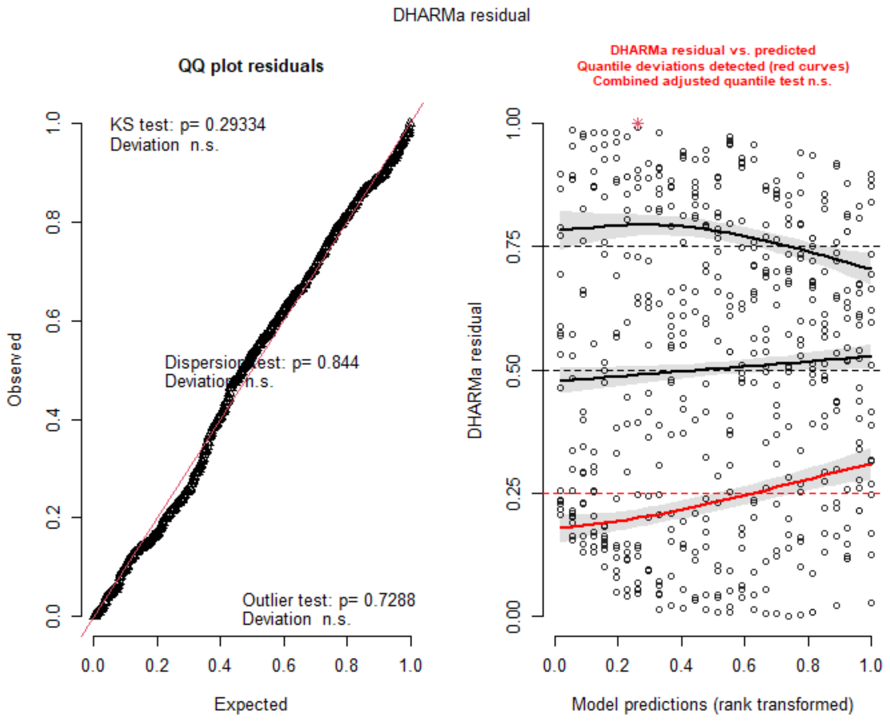

### R. Cocaine Infusions First 15 Minutes by Pre-Lever Activity

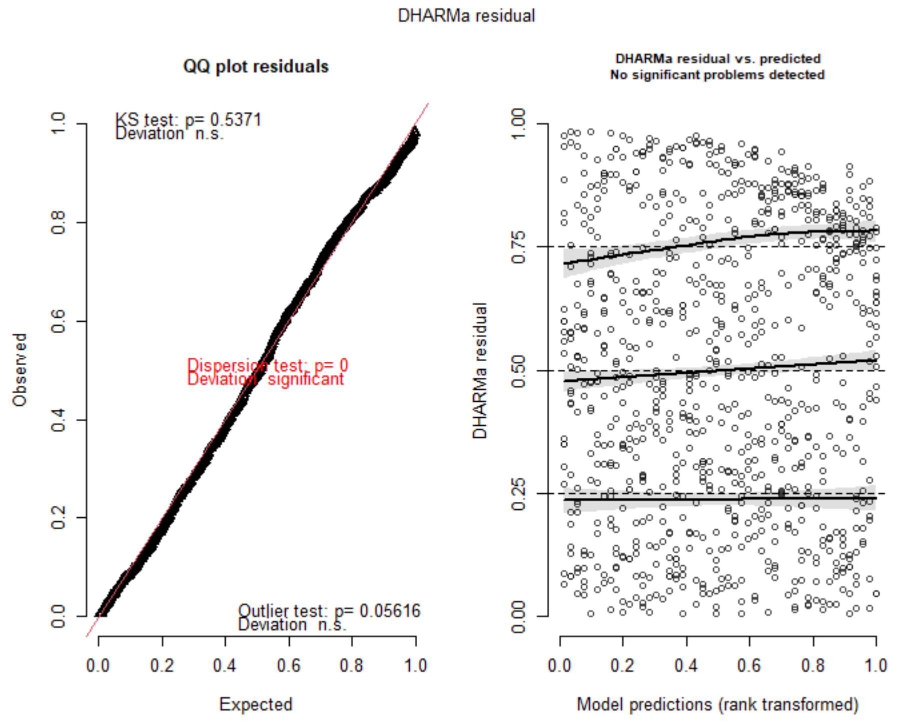


### S. Nose Motion Percent Difference From Drug 01 by Behavioral Expression

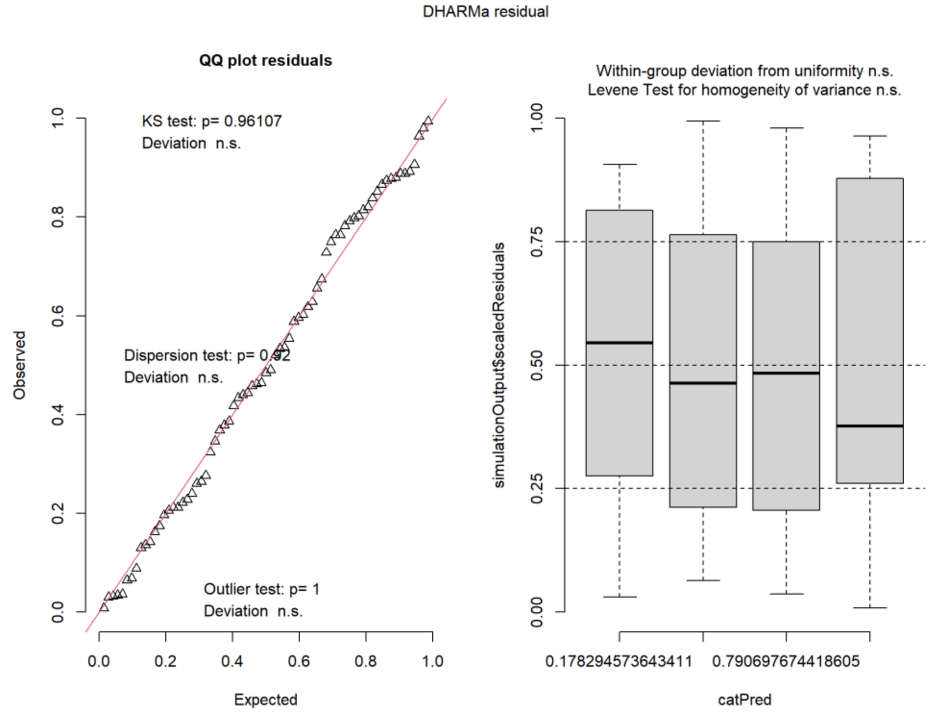

### T. Cocaine Infusions in Short Access by Sex

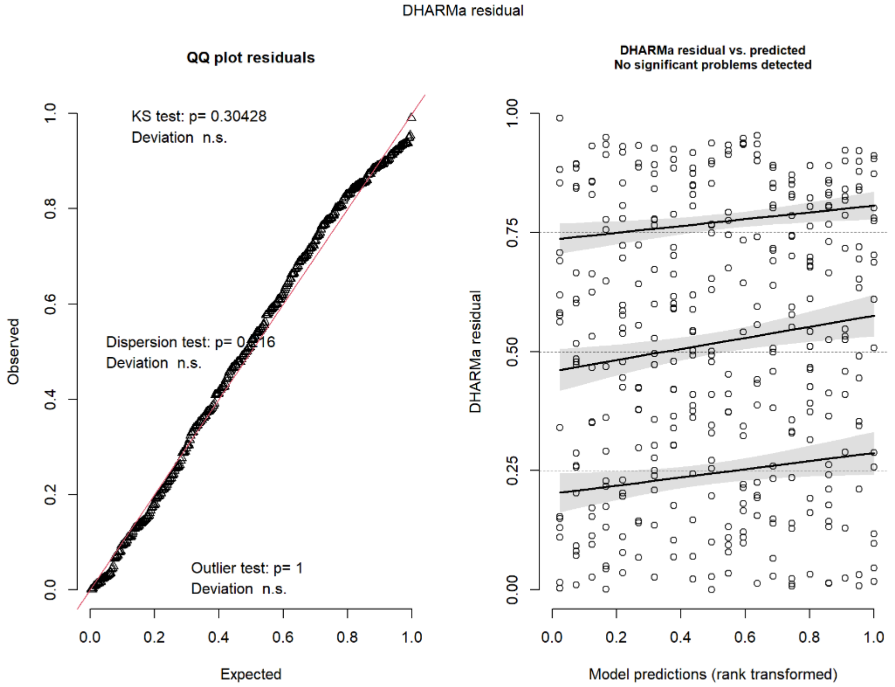


### U. Cocaine Infusions in Long Access by Sex

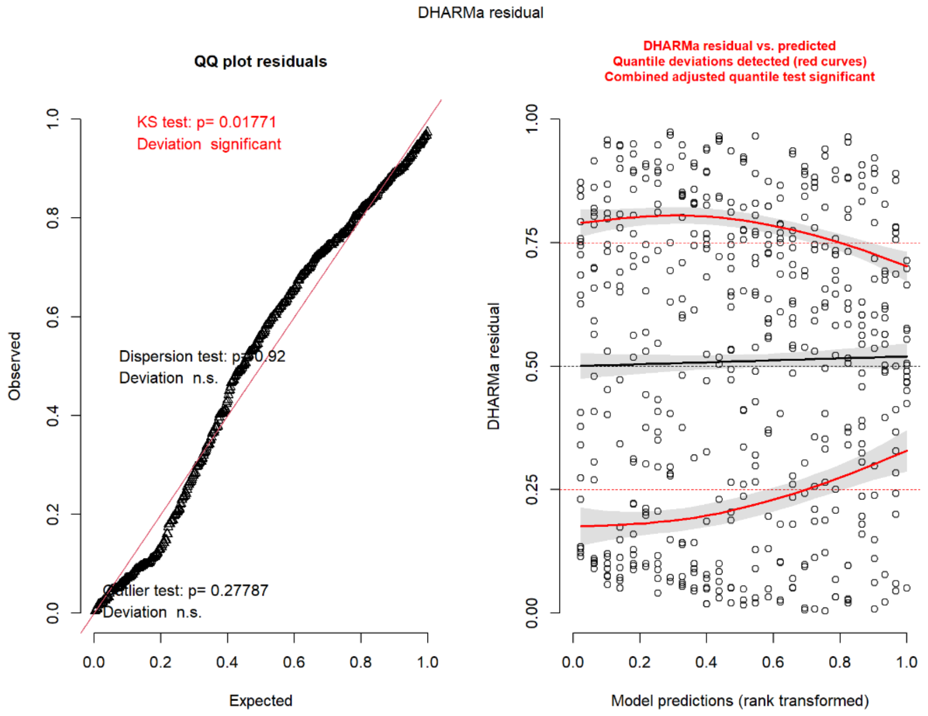

### V. Locomotion Percent Difference From Baseline in Noncontingent Sessions by Sex

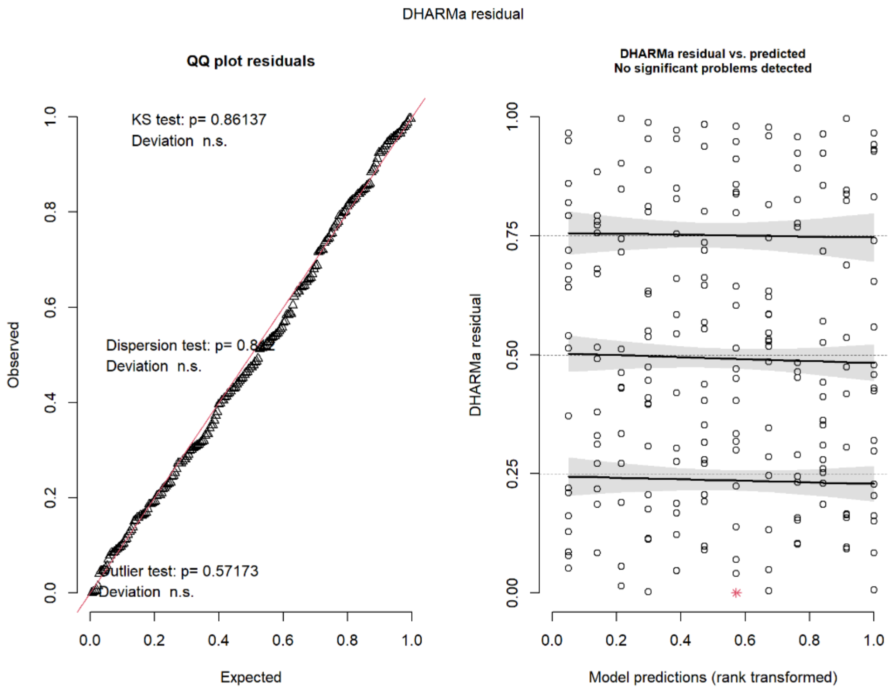


### W. Cocaine Infusions in Short Access by Behavioral Expression and Sex

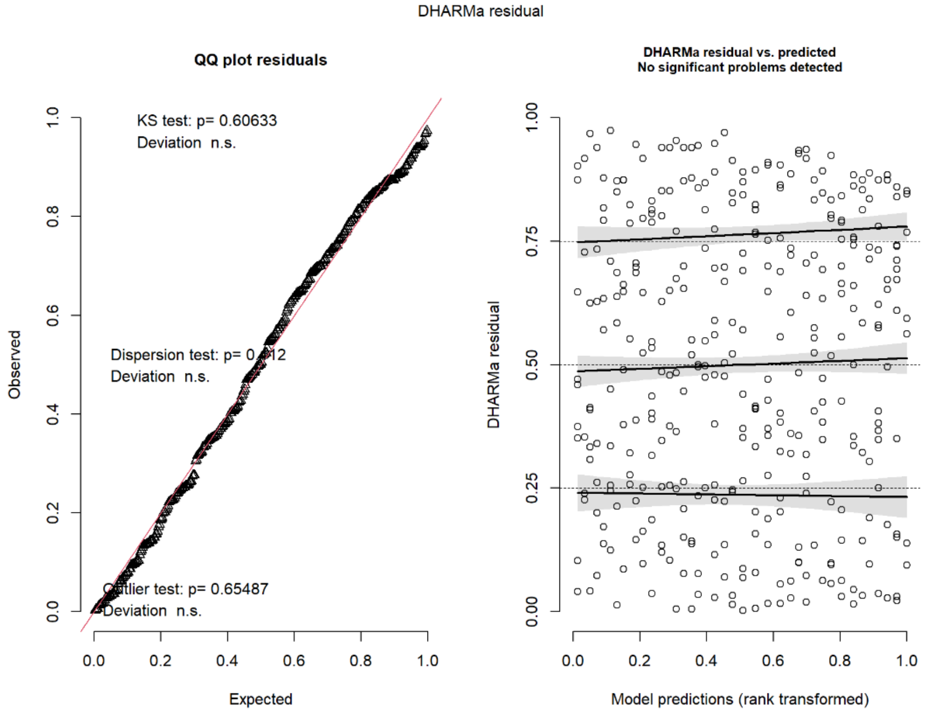

### X. Cocaine Infusions in Long Access by Behavioral Expression by Sex

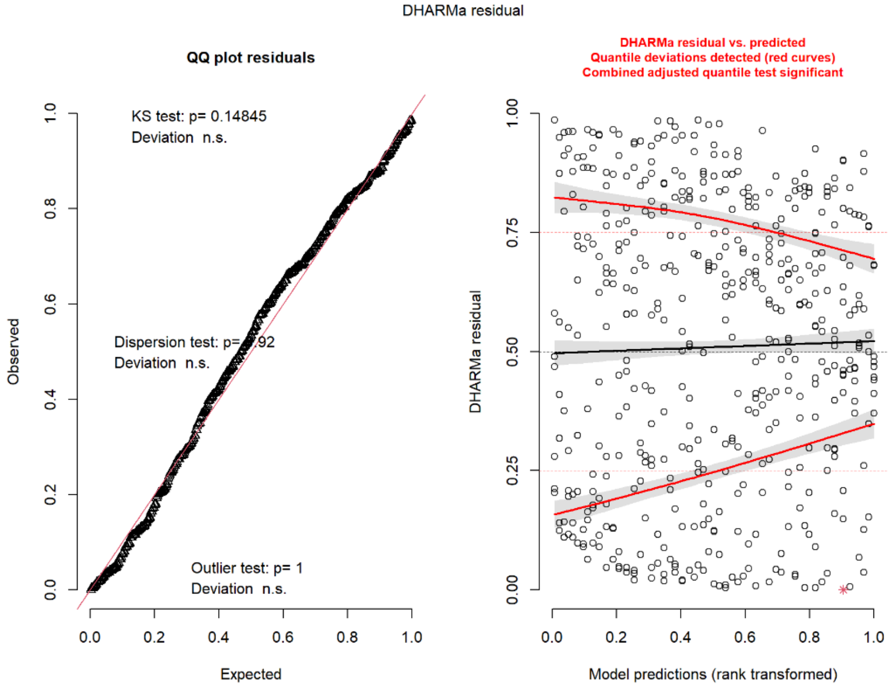


### Y. Active Lever Entrances per Meter by Session Type, Abstinence, and Sex

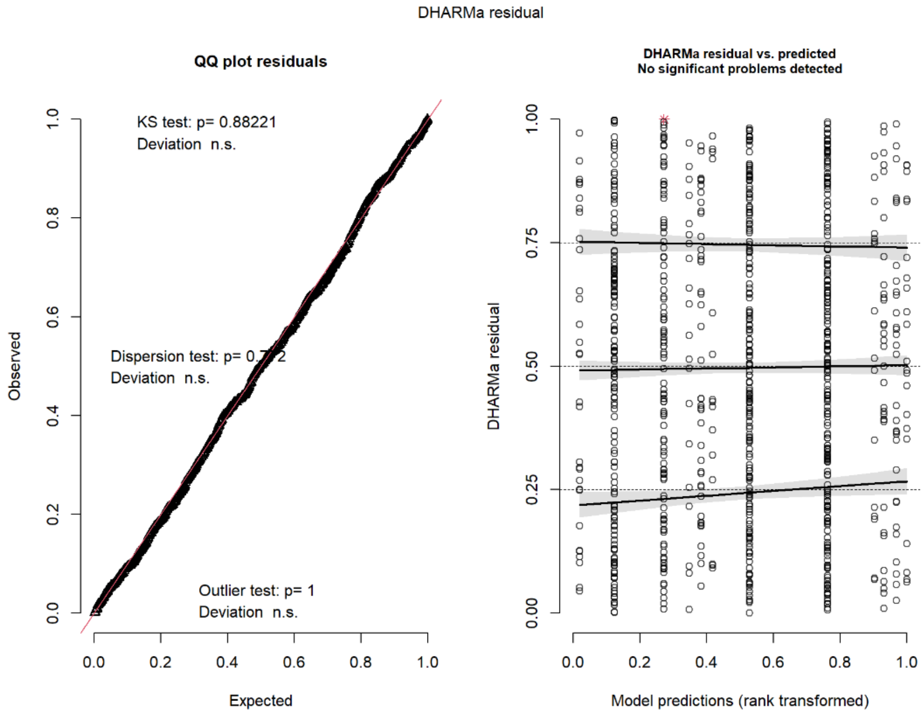

### Z. Entrances per Meter Percent Difference From Noncontingent 01 by Sex

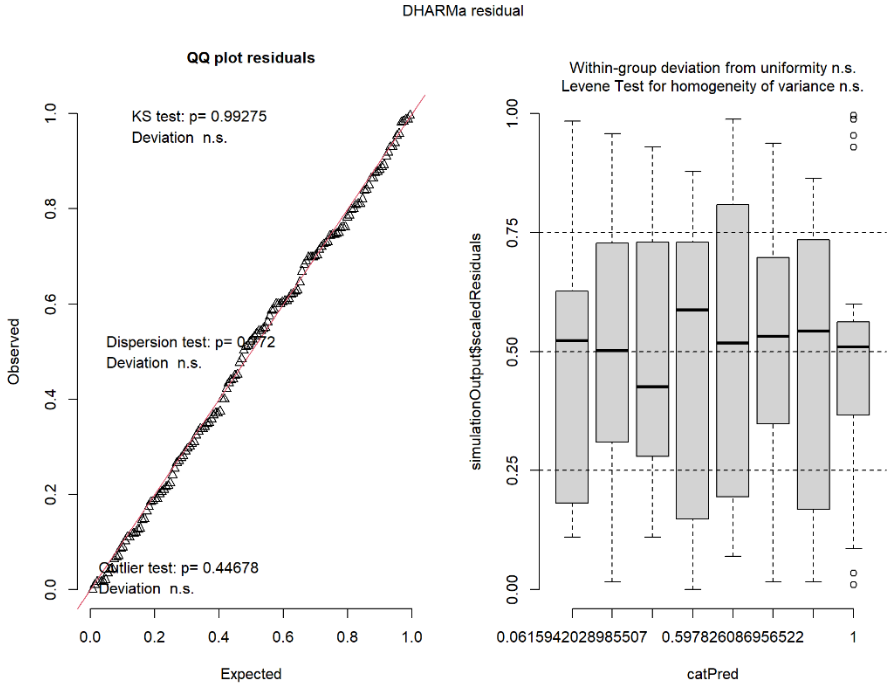


### AA. Cocaine Infusions in Short Access by Pre-Lever Activity and Sex

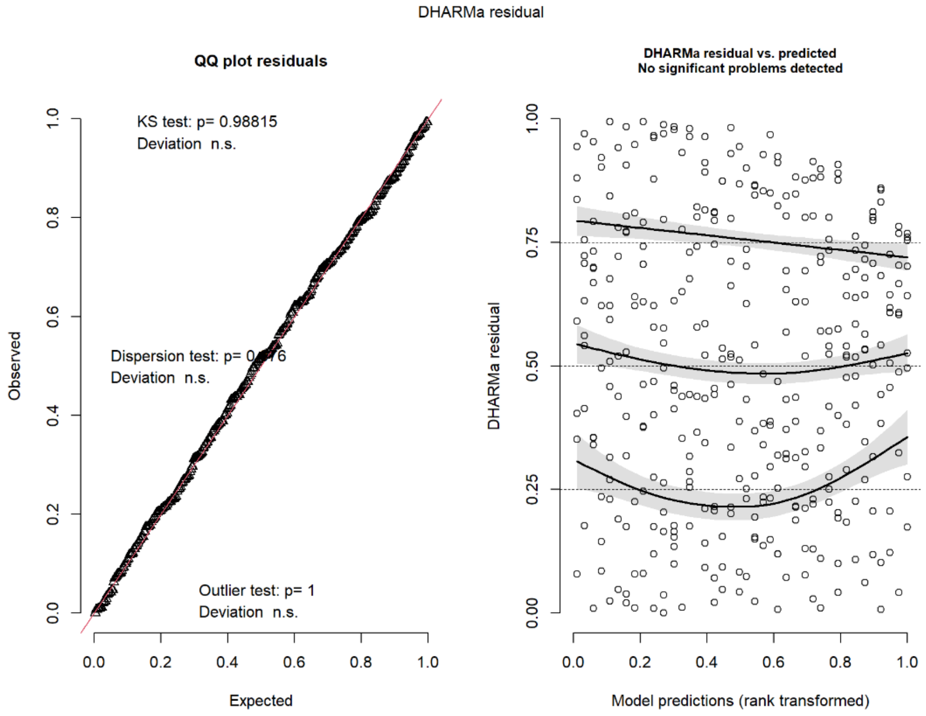

**BB. Cocaine Infusions in Long Access by Pre-Lever Activity and Sex**

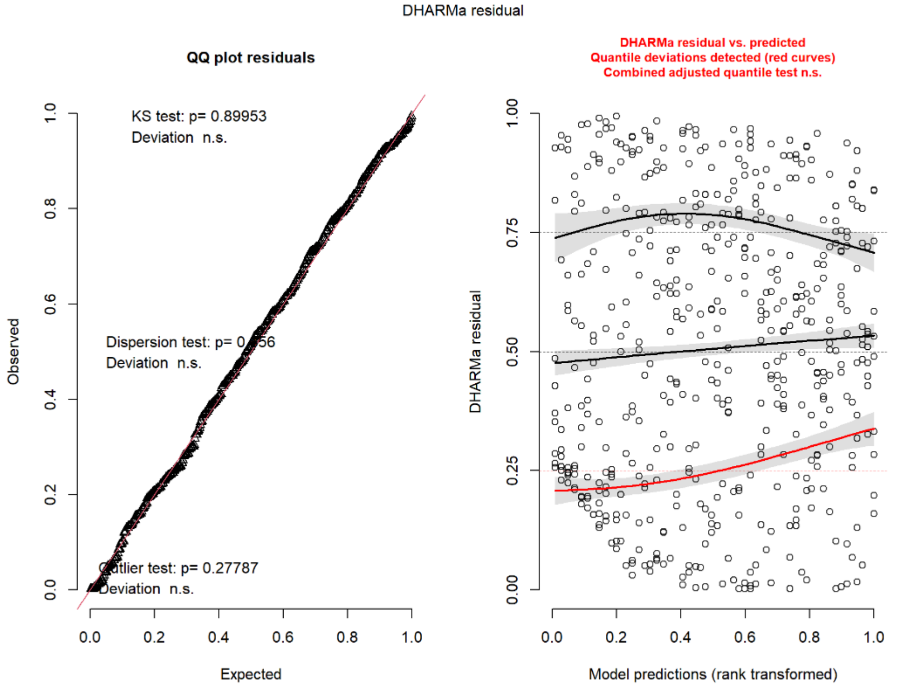


### 3. Nose Motion Analysis

#### 3.1. Tolerant rats show no evidence of stereotypies during Drug sessions

To assess if rats in the Tolerance group were miscategorized due to stereotypies, nose motion percentage change from Baseline (Drug 01) between behavioral expression groups was analyzed. Following a 0.0002 lambda Box-Cox power transformation, a Type III Wald test found a significant main effect of behavioral expression ($\chi^2(1)=5.77$, $p=.034$), of session ($\chi^2(1)=9.87$, $p=.005$) and their interaction ($\chi^2(1)=18.95$, $p<.001$). Post hoc analyses revealed rats in the Tolerance group to have significantly decreased their nose motion in Drug 03 compared to both baseline and Drug 02. Conversely, the Sensitization group showed significantly increased nose motion in both Drug 02 and Drug 03 compared to baseline, and Drug 03 greater than

Drug 02. Comparatively, the Sensitization group had greater nose motion within Drug 02 and Drug 03 than the Tolerance group (**Fig. S1**). These results demonstrate that the observed effect seen in **Fig. 2C** was not due to stereotypies, and that behavioral expression groups were sufficiently categorized based on their locomotor difference from Drug 01 to Drug 03.

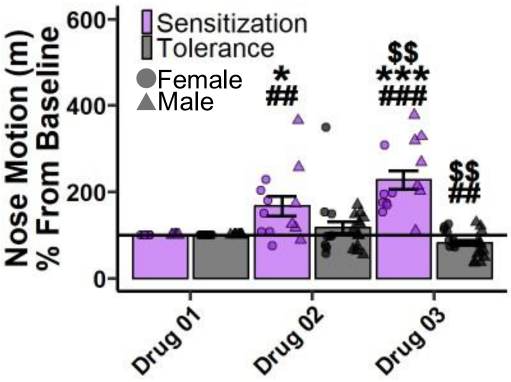


**Supplementary Figure 1. Tolerant rats show no evidence of stereotypies during Drug sessions.** Nose motion (m) percentage difference from Baseline (Drug 01) grouped by psychomotor behavioral expression. ##*p*<.01, ###*p*<.001 vs Baseline*;* Within behavioral expression group: $$*p*<.01 vs Drug 02; Between behavioral expression groups: **p*<.05 vs Tolerance Drug 02; ****p*<.001 vs Tolerance Drug 03.

### 4. Sex as a Factor Analyses

The following analyses were conducted with sex included as a factor. These specific tests were chosen as they are critical results to the manuscript interpretation. The main focus was the full interaction between factors and reported below.

#### 4.1. No effect of sex on cocaine infusions in Short Access

To determine if sexes differed in cocaine infusions during Short Access, a Type III Wald test was performed. Results showed no significant interaction between sex and session ($\chi^2(9)=14.3$, $p=.233$). These results show that cocaine intake did not differ between sexes in Short Access.

#### 4.2. No effect of sex on cocaine infusions in Long Access

To determine if sexes differed in cocaine infusions during Long Access, a Type III Wald test was performed. Results showed no significant interaction between sex and session ($\chi^2(13)=16.77$, $p=.583$). These results show that cocaine intake did not differ between sexes in Long Access.

#### 4.3. No effect of sex on locomotion percent difference from baseline during Noncontingent sessions

To determine if sexes differed in their locomotion perfect difference from baseline during Noncontingent sessions, a Type III Wald test was performed. Results showed no significant interaction between sex and session ($\chi^2(5)=10.72$, $p=.159$). These results show that locomotion percent difference from baseline did not differ between sexes in Noncontingent sessions.

#### 4.4. No effect of sex x behavioral expression on cocaine infusions across Short Access sessions

To determine if a sex x behavioral expression x session interaction had an effect on cocaine infusions during Short Access, a Type III Wald test was performed. Results showed the interaction was not significant ($\chi^2(9)=14.31$, $p=.607$), indicating that cocaine intake did not differ between sexes as a function of behavioral expression across Short Access sessions.

#### 4.5. No effect of sex x behavioral expression on cocaine infusions across Long Access sessions

To determine if a sex x behavioral expression x session interaction had an effect on cocaine infusions during Long Access, a Type III Wald test was performed. Results showed no significant interaction effect ($\chi^2(13)=9.48$, $p=.736$), indicating that cocaine intake did not differ between sexes as a function of behavioral expression across Long Access sessions.

#### 4.6. No effect of sex x session type x abstinence on active lever entrances per meter pre-lever

To determine if a sex x session type x abstinence interaction had an effect on active lever entrances per meter pre-lever, a Type III Wald test was performed. Results showed no significant interaction effect ($\chi^2(2)=3.96$, $p=.138$), indicating that active lever entrances per meter did not differ between sexes as a function of session type and abstinence.

#### 4.7. No effect of sex x levers on entrances per meter percent difference from Noncontingent 01

To determine if a sex x levers interaction had an effect on entrances per meter perfect difference from Noncontingent 01, a Type III Wald test was performed. Results showed no significant interaction effect ($\chi^2(1)=0.05$, $p=1$), indicating that entrances per meter percent difference from Noncontingent 01 did not differ between sexes towards either the active or inactive lever during Noncontingent sessions.

#### 4.8. No effect of sex x pre-lever activity group on cocaine infusions across Short Access

To determine if a sex x pre-lever activity group interaction had an effect on cocaine infusions during Short Access, a Type III Wald test was performed. Results showed no significant interaction between sex and pre-lever activity groups ($\chi^2(9)=15.61$, $p=.438$). These results show that cocaine intake did not differ between sexes as a function of pre-lever activity group across Short Access sessions.

#### 4.9. An effect of sex x pre-lever activity group observed on cocaine infusions across Long Access

To determine if a sex x pre-lever activity group interaction had an effect on cocaine infusions during Long Access, a Type III Wald test was performed. Results showed a significant interaction between sex x pre-lever activity group x session ($F(13,467.25)=2.34$, $p=.023$). Post hoc analyses revealed Low Pre-Lever Activity females to have greater cocaine infusions than Low Pre-Lever Activity males from Long Access 04 to Long Access 09, excluding Long Access 07, suggesting that females are more sensitive to escalation of cocaine intake (**Fig. S2**). These results align with previous reports that females escalate in use before males (de Guglielmo et al., 2024).

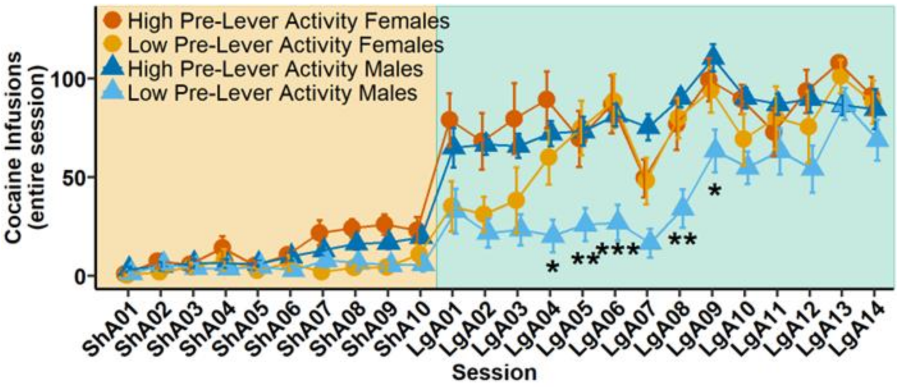


**Supplementary Figure 2. Low Pre-Lever Activity Female rats receive greater cocaine infusions than Low Pre-Lever Activity Male rats during Long Access.**

Cocaine infusions across Short and Long Access by sex and pre-lever activity group. Within Low Pre-Lever Activity group: **p*<.05, ***p*<.01, ****p*<.001 vs Females.

behaviors reveals that escalation of intake, aversion-resistant responding, and breaking-points are highly correlated measures of the same construct. *eLife*, *12*, RP90422. https://doi.org/10.7554/eLife.90422
