## Supplementary material for "Incentive Salience, not Psychomotor Sensitization or Tolerance, Drives Escalation of Cocaine Self-Administration in Heterogeneous Stock Rats": Final Mixed Models for Analyses

**Mixed Effects Models Used in Analyses**

| Manuscript Plot | Mixed Model Type | Response Variable | Fixed Effects | Random Effects | | | | Dispersion Modeling | Distribution Family (link) | AIC | BIC | Log-Likelihood | Deviance | Observations | DHARMa Plot |
|---|---|---|---|---|---|---|---|---|---|---|---|---|---|---|---|
| | | | | Group | Effect | Std. Dev. | Variance | | | | | | | | |
| Figure 1.B. | Generalized | Cocaine Infusions Short Access | Session | Rat | Intercept | 1.42 | 1.19 | Abstinence | nbinomal1 (log) | 1940.9 | 1995.3 | -956.4 | 1912.9 | 362 | A |
| Figure 1.B. | Generalized | Cocaine Infusions Long Access | Session | Rat | Intercept | 568.4 | 23.84 | N/A | gaussian (identity) | 4822 | 4889.6 | -2395 | 4790 | 504 | B |
| | | | | Residual | - | 651 | 25.51 | | | | | | | | |
| Figure 1.C. | Generalized | Cocaine Infusions First 15mins | Session | Rat | Intercept | 0.59 | 0.77 | Session | tweedie (log) | 2669.6 | 2907.8 | -1284.8 | 2569.6 | 866 | C |
| Figure 2.A. | Linear | Locomotion Raw Noncontingent Sessions | Session | Rat | Intercept | 0.09 | 0.29 | N/A | gaussian (identity) | 1140.9 | 1182.6 | -559.5 | 1118.9 | 327 | D |
| | | | | Residual | - | 1.72 | 1.31 | | | | | | | | |
| Figure 2.B. | Linear | Locomotion Percent Difference From Baseline Noncontingent Sessions | Session | Rat | Intercept | 0.1 | 0.31 | N/A | gaussian (identity) | 369.6 | 396.4 | -176.8 | 353.6 | 212 | E |
| | | | | Residual | - | 0.26 | 0.51 | | | | | | | | |
| Figure 2.C. | Generalized | Locomotion Percent Difference From Drug 01 | Session * Behavioral Expression | Rat | Intercept | 0.13 | 0.35 | Behavioral Expression | gaussian (identity) | 187.4 | 203.2 | -86.7 | 173.4 | 71 | F |
| Figure 2.D. | Generalized | Cocaine Infusions Short Access | Session * Behavioral Expression | Rat | Intercept | 1.46 | 1.21 | Session | tweedie (log) | 1966.9 | 2090.5 | -951.4 | 1902.9 | 352 | G |
| Figure 2.D. | Generalized | Cocaine Infusions Long Access | Session * Behavioral Expression | Rat | Intercept | 580.6 | 24.1 | Session * Behavioral Expression | gaussian (identity) | 4706.2 | 4945.3 | -2296.1 | 4592.2 | 490 | H |
| Figure 2.E | Generalized | Cocaine Infusions First 15mins | Session * Behavioral Expression | Rat (Conditional) | Intercept | 0.05 | 0.23 | Session * Behavioral Expression + Rat | nbinomial12 (log) | 2733.3 | 3202.2 | -1267.7 | 2535.3 | 842 | I |
| | | | | Rat (Dispersion) | Intercept | 7.3 | 2.7 | | | | | | | | |
| Figure 3.A | Generalized | Locomotion Pre-Lever | Session | Rat | Intercept | 0.34 | 0.58 | Abstinence | gaussian (identity) | 2877 | 3010.2 | -1410.5 | 2821 | 860 | J |
| Figure 3.B. | Generalized | Locomotion Pre-Lever | Session Type * Abstinence | Rat | Intercept | 0.32 | 0.57 | Session Type * Abstinence* Infusion Group | gaussian (identity) | 2875 | 2965.3 | -1418.5 | 2837 | 860 | K |
| Figure 3.H | Generalized | Active Lever Entrances per Meter Pre-Lever | Session | Rat (Conditional) | Intercept | 0.15 | 0.39 | Session Type * Abstinence + Rat | tweedie (log) | 1475.3 | 1632.2 | -704.6 | 1409.3 | 859 | L |
| | | | | Rat (Dispersion) | Intercept | 0.11 | 0.34 | | | | | | | | |
| Figure 3.I. | Generalized | Active Lever Entrances per Meter Pre-Lever | Session Type * Abstinence | Rat (Conditional) | Intercept | 0.14 | 0.38 | Session + Rat | tweedie (log) | 1514.7 | 1671.6 | -724.3 | 1448.7 | 859 | M |
| | | | | Rat (Dispersion) | Intercept | 0.13 | 0.37 | | | | | | | | |
| Figure 4.A. | Generalized | Lever Entrances per Meter Percent Difference From Noncontingent 01 | Session | Rat (Conditional) | Intercept | 0.83 | 0.91 | Session + Rat | gaussian (identity) | 731.1 | 755.1 | -357.5 | 715.1 | 148 | N |
| | | | | Rat (Dispersion) | Intercept | 0.17 | 0.42 | | | | | | | | |
| Figure 4.C. | Generalized | Active Lever Entrances per Meter Percent Difference From Noncontingnet 01 | Session * Pre-Lever Activity | Rat (Conditional) | Intercept | 0.11 | 0.33 | Pre-Lever Activity + Rat | gaussian (identity) | 361.5 | 379.9 | -172.8 | 345.5 | 74 | O |
| | | | | Rat (Dispersion) | Intercept | 0.54 | 0.74 | | | | | | | | |
| Figure 4.D. | Generalized | Cocaine Infusions Short Access | Session * Pre-Lever Activity | Rat | Intercept | 1.27 | 1.13 | Abstinence | tweedie (log) | 1947.1 | 2044.4 | -948.6 | 1897.1 | 362 | P |
| Figure 4.D. | Generalized | Cocaine Infusions Long Access | Session * Pre-Lever Activity | Rat | Intercept | 364.6 | 19.1 | Pre-Lever Activity | gaussian (identity) | 4807.3 | 4938.2 | -2372.6 | 4745.3 | 504 | Q |
| Figure 4.E. | Generalized | Cocaine Infusions First 15mins | Session * Pre-Lever Activity | Rat (Conditional) | Intercept | 0.09 | 0.3 | Session + Rat | nbinomial12 (log) | 2737 | 3094.3 | -1293.5 | 2587 | 866 | R |
| | | | | Rat (Dispersion) | Intercept | 3.81 | 1.95 | | | | | | | | |
| Supplementary Figure 1. | Generalized | Nose Motion Percent Difference From Drug 01 | Session * Behavioral Expression | Rat | Intercept | 0.05 | 0.22 | Session * Behavioral Expression | gaussian (identity) | 87.2 | 109.9 | -33.6 | 67.2 | 61 | S |
| | | | | Rat (Dispersion) | Intercept | 3.26E-10 | 1.81E-05 | | | | | | | | |
| N/A | Generalized | Cocaine Infusions Short Access | Session * Sex | Rat | Intercept | 1.45 | 1.21 | Session | nbinomal1 (log) | 1952.6 | 2073.2 | -945.3 | 1890.6 | 331 | T |
| N/A | Generalized | Cocaine Infusions Long Access | Session * Sex | Rat | Intercept | 527.1 | 22.96 | Session | gaussian (identity) | 4829.1 | 5010.7 | -2371.6 | 4743.1 | 461 | U |
| N/A | Generalized | Locomotion Percent Difference From Baseline Noncontingent Sessions | Session * Sex | Rat | Intercept | 0.1 | 0.31 | Sex | gaussian (identity) | 372.6 | 423 | -171.3 | 342.6 | 197 | V |
| N/A | Generalized | Cocaine Infusions Short Access | Session * Behavioral Expression * Sex | Rat | Intercept | 1.53 | 1.24 | Session * Sex | tweedie (log) | 1978.7 | 2218.3 | -927.4 | 1854.7 | 290 | W |
| N/A | Generalized | Cocaine Infusions Long Access | Session * Behavioral Expression * Sex | Rat | Intercept | 422.6 | 20.56 | Abstinence * Sex | gaussian (identity) | 4715.8 | 4980.1 | -2294.9 | 4589.8 | 427 | X |
| N/A | Generalized | Active Lever Entrances per Meter Pre-Lever | Session Type * Abstinence * Sex | Rat (Conditional) | Intercept | 0.13 | 0.37 | Session * Sex + Rat | tweedie (log) | 1532.3 | 1831.9 | -703.2 | 1406.3 | 796 | Y |
| | | | | Rat (Dispersion) | Intercept | 0.14 | 0.37 | | | | | | | | |
| N/A | Generalized | Lever Entrances per Meter Percent Difference From Noncontingent 01 | Session * Levers * Sex | Rat (Conditional) | Intercept | 1.02 | 1.01 | Session * Sex + Rat | gaussian (identity) | 744.9 | 786.9 | -358.5 | 716.9 | 134 | Z |
| | | | | Rat (Dispersion) | Intercept | 0.11 | 0.33 | | | | | | | | |
| Supplementary Figure 2. | Generalized | Cocaine Infusions Short Access | Session * Pre-Lever Activity * Sex | Rat | Intercept | 1.27 | 1.13 | Pre-Lever Activity * Sex | tweedie (log) | 1944.1 | 2123.2 | -926.1 | 1852.1 | 316 | AA |
| Supplementary Figure 2. | Linear | Cocaine Infusions Long Access | Session * Pre-Lever Activity * Sex | Rat | Intercept | 290.2 | 17.04 | N/A | gaussian (identity) | 4811.3 | 5056.2 | -2347.6 | 4695.3 | 446 | BB |
| | | | | Residual | - | 558.3 | 23.63 | | | | | | | | |
